# αKG-mediated carnitine synthesis promotes homologous recombination via histone acetylation

**DOI:** 10.1101/2024.02.06.578742

**Authors:** Apoorva Uboveja, Zhentai Huang, Raquel Buj, Amandine Amalric, Hui Wang, Naveen Kumar Tangudu, Aidan R. Cole, Emily Megill, Daniel Kantner, Adam Chatoff, Hafsah Ahmad, Mariola M. Marcinkiewicz, Julie A. Disharoon, Sarah Graff, Erika S. Dahl, Nadine Hempel, Wayne Stallaert, Simone Sidoli, Benjamin G. Bitler, David T. Long, Nathaniel W. Snyder, Katherine M. Aird

## Abstract

Homologous recombination (HR) deficiency enhances sensitivity to DNA damaging agents commonly used to treat cancer. In HR-proficient cancers, metabolic mechanisms driving response or resistance to DNA damaging agents remain unclear. Here we identified that depletion of alpha-ketoglutarate (αKG) sensitizes HR-proficient cells to DNA damaging agents by metabolic regulation of histone acetylation. αKG is required for the activity of αKG-dependent dioxygenases (αKGDDs), and prior work has shown that changes in αKGDD affect demethylases. Using a targeted CRISPR knockout library consisting of 64 αKGDDs, we discovered that Trimethyllysine Hydroxylase Epsilon (TMLHE), the first and rate-limiting enzyme in *de novo* carnitine synthesis, is necessary for proliferation of HR-proficient cells in the presence of DNA damaging agents. Unexpectedly, αKG-mediated TMLHE-dependent carnitine synthesis was required for histone acetylation, while histone methylation was affected but dispensable. The increase in histone acetylation via αKG-dependent carnitine synthesis promoted HR-mediated DNA repair through site- and substrate-specific histone acetylation. These data demonstrate for the first time that HR-proficiency is mediated through αKG directly influencing histone acetylation via carnitine synthesis and provide a metabolic avenue to induce HR-deficiency and sensitivity to DNA damaging agents.

## Introduction

The ability of cells to accurately and efficiently repair DNA damage, and in particular DNA double strand breaks (DSBs), is critical for genome stability (van Gent et al., 2001). In cancer, DNA repair pathways are often dysregulated (Brown et al., 2017; Groelly et al., 2023; Helleday et al., 2008). Homologous recombination (HR), an error-free DNA DSB repair mechanism, is deficient in many tumors, leading to enhanced sensitivity to widely clinically used DNA damaging agents. Interestingly, HR-proficient tumors are often harder to treat due to their intrinsic resistance to DNA damaging agents (Konstantinopoulos et al., 2015). For instance, tumors with high endogenous *CCNE1* expression, which encodes the oncogene cyclin E1, are HR-proficient and inherently resistant to DNA damaging agents (Patch et al., 2015). Mechanisms underlying HR-proficiency in general, and in the context of CCNE1, remain unclear.

Epigenetic modifications such as histone tail post-translational modifications (PTMs) play a crucial role in many fundamental cellular processes including DNA DSB repair (Gong and Miller, 2013, 2019; Song et al., 2023). Interestingly, histone PTMs are metabolically sensitive (Izzo et al., 2021). For instance, alpha-ketoglutarate (αKG), also known as 2-oxoglutarate (2OG), is a co-substrate for αKG-dependent dioxygenases (αKGDDs) that include DNA and histone demethylases (Islam et al., 2018). In humans, there are >60 different αKGDDs, which are a superfamily of enzymes that reside at the intersection of cancer metabolism and cancer epigenetics and play crucial roles in many biological processes. αKGDDs catalyze hydroxylation reactions on a variety of substrates and require the presence of αKG, Fe[II], and oxygen, producing succinate. Alteration of several metabolic enzymes and pathways that affect αKG abundance and αKGDD activity have been reported in cancer, such as mutations of IDH1 and IDH2, changes in glutaminolysis that deplete αKG, or alterations in fumarate hydratase (FH) and succinate dehydrogenase (SDH) that inhibit αKGDD activity (Losman et al., 2020). Prior work has found that oncometabolites that suppress αKGDD activity affect DNA repair and sensitivity to DNA damaging agents (Inoue et al., 2016; Jiang et al., 2015; Molenaar et al., 2018; Sulkowski et al., 2017; Sulkowski et al., 2020; Sulkowski et al., 2018; Wang et al., 2015). Thus far, the majority of studies on αKG and αKGDDs have focused on the handful of DNA and histone methylases. However, since there are >60 αKGDDs, there is a strong likelihood that changes in αKG abundance affects DNA repair processes in multiple ways.

Prior work has demonstrated that histone acetylation is necessary for HR-mediated DNA repair (Gong and Miller, 2013; Song et al., 2023). Carnitine functions as a carrier for acyl-groups into and out of the mitochondria (Longo et al., 2016), and acetylcarnitine shuttling from the mitochondria and peroxisome can provide the acetyl groups for histone acetylation (Izzo et al., 2023; Kuna et al., 2023). Trimethyllysine Hydroxylase Epsilon (TMLHE), an αKGDD, is the first and rate-limiting step in *de novo* carnitine synthesis but no studies to date have characterized how changes in αKG affect carnitine synthesis or carnitine-dependent processes.

Here, we found that depletion of αKG sensitized HR-proficient cells to DNA damaging agents. Using a CRISPR KO screen targeting 64 αKGDDs, we identified and validated TMLHE as the aKGDD driving these effects. αKG promoted carnitine synthesis and carnitine-mediated histone acetylation, whereas histone methylation was less robustly affected. We identified 3 specific histone acetyl marks that are regulated in an αKG- and carnitine-dependent manner (H4K8ac, H4K12ac, and K3K23ac) and found that the association of these marks at DSBs are decreased by depletion of αKG and TMLHE and rescued by carnitine. Moreover, this axis is critical for HR-mediated DNA repair. Together, these studies identified a histone acetylation pathway that is promoted by αKG production and suggest demethylation reactions are dispensable for the observed HR-proficiency and corresponding resistance to DNA damaging agents.

## Results

### The αKG-dependent carnitine synthesis enzyme TMLHE is required for CCNE1-driven cell proliferation in the presence of DNA damaging agents

We previously published that wildtype IDH1 and αKG are increased in high grade serous ovarian cancer (HGSOC) cell lines compared to normal fallopian tube cells (Dahl et al., 2019). Upon further molecular analysis, we found that the HGSOC cell lines used have amplification or overexpression of *CCNE1* (encoding cyclin E). Thus, we aimed to determine whether CCNE1 itself drives αKG abundance. Towards this goal, we overexpressed CCNE1 in two ovarian cancer-relevant cell models (**Fig. S1A**). We found that overexpression of CCNE1 increased αKG abundance in both isogenic cell lines (**Fig. 1A**). αKG is derived from multiple metabolic pathways, including oxidative decarboxylation of isocitrate via isocitrate dehydrogenases (IDHs) and glutaminolysis (**Fig. 1B**). We found that depletion of αKG using either an IDH1 inhibitor (**Fig. 1A**) or glutamine starvation (**Fig. S1B**) sensitized CCNE1-driven cells to the DNA damaging agents olaparib [a poly(ADP-ribose) polymerase (PARP) inhibitor] and cisplatin (**Fig. 1C-D**). Inhibition of IDH1 also sensitized CCNE1-driven tumors to olaparib *in vivo* (**Fig. 1E**). Inhibition of IDH1 did not sensitize CCNE1-driven cells to the microtubule stabilizing agent paclitaxel (**Fig. S1**), suggesting the effect is specific for DNA damaging agents. The IDH1 inhibitor used was developed against mutant IDH1 but targets the wildtype enzyme at higher concentrations (Calvert et al., 2017; Dahl et al., 2019), and wildtype IDH1 overexpression fully rescued the phenotype, suggesting the observations are not due to off-target effects (**Fig. S1D-E**). While the IDH1 inhibitor decreased both αKG and citrate abundance (**Fig. 1A and S1F**), only supplementing with αKG but not citrate rescued the effects (**Fig. 1F)**. Similarly, glutamine starvation was rescued by αKG supplementation (**Fig. 1G**). αKG is required as a co-substrate for αKGDDs in mammalian cells, and αKGDDs are inhibited by succinate (Islam et al., 2018). Interestingly, succinate supplementation also phenocopied decreased αKG in its ability to inhibit proliferation of CCNE1-driven cells in the presence of DNA damaging agents and was rescued by αKG supplementation (**Fig. S1G**), suggesting this effect is due to decreased activity of αKGDDs. To ascertain which αKGDD is driving the observed phenotype, we constructed a CRISPR knockout library of 64 αKGDDs (**Table S1**) and performed a dropout screen in the presence of olaparib (**Fig. 1H**). Unexpectedly, TMLHE, the first and rate-limiting enzyme *in de novo* carnitine synthesis, was the only one of the top five genes to drop out in both CCNE1-driven cell lines (**Fig. 1I-J**). Using TMLHE shRNA (**Fig. S1H-I**) and the carnitine synthesis inhibitor mildronate, we validated this observation in both cell lines using both olaparib and cisplatin (**Fig. 1K-L and S1J-K)**. Supplementing TMLHE knockdown cells with L-carnitine, but not αKG, rescued the proliferation defects in the presence of DNA damaging agents, confirming that αKG was upstream of TMLHE-catalyzed carnitine synthesis (**Fig. 1K-L and S1J-K**). Similarly, supplementation of L-carnitine rescued the loss of proliferation observed in cells treated with olaparib or cisplatin in combination with αKG-depleting conditions or succinate (**Fig. 1M-N and S1L-N**). Together, these data place carnitine downstream of αKG and demonstrate the necessity of the αKGDD TMLHE and *de novo* carnitine synthesis for proliferation in response to DNA damaging agents in CCNE1-driven, HR-proficient models (**Fig. 1O**).

**Figure 1.**
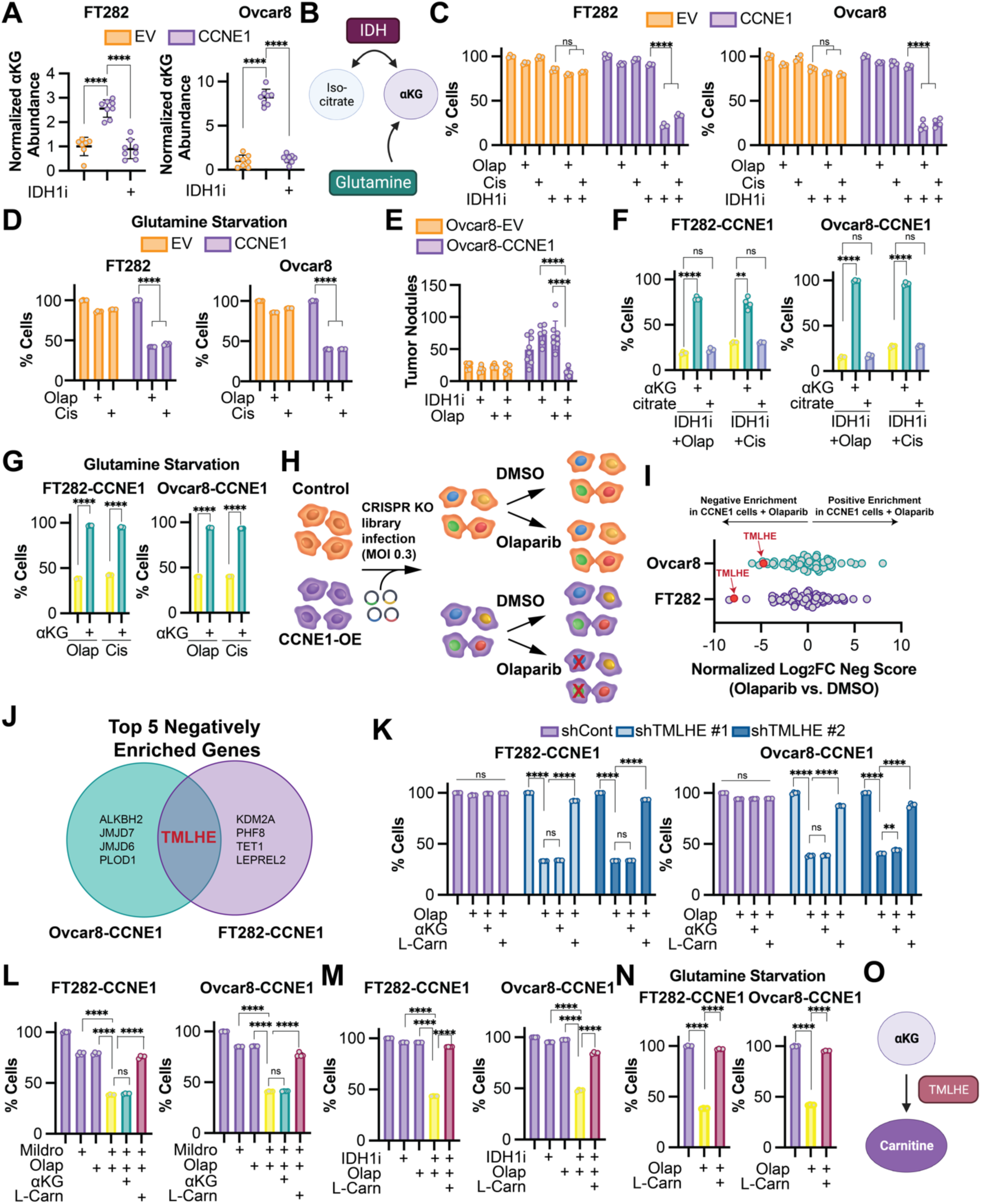
CRISPR drop out screen identifies the αKG-dependent dioxygenase TMLHE as a requirement for CCNE1-driven cell proliferation in response to DNA damaging agents. **A)** The indicated cells were treated with the IDH1 inhibitor GSK864 (IDH1i), and αKG abundance was assessed by LC-MS. Cells expressing empty vector (EV) = orange; Cells expressing CCNE1 (CCNE1) = purple. **B)** Schematic of αKG synthesis relevant to this project. **C)** The indicated CCNE1-low (orange) and -high (purple) isogenic cells were treated with the IDH1 inhibitor GSK864 (IDH1i) and DNA damaging agents olaparib (Olap) or cisplatin (Cis) alone or in combination. % of cells was assessed by crystal violet staining and normalized to controls. **D)** The indicated CCNE1-low (orange) and -high (purple) isogenic cells were cultured under glutamine starvation conditions and treated with DNA damaging agents olaparib (Olap) or cisplatin (Cis) alone or in combination. % of cells was assessed by crystal violet staining and normalized to vehicle controls. **E)** Ovcar8 isogenic cells were injected IP into immunocompromised female mice (n=8/group). Cells expressing empty vector (EV) = orange; Cells expressing CCNE1 (CCNE1) = purple. The mice were treated with vehicle, the IDH1 inhibitor GSK864 (IDH1i), and olaparib (Olap) alone or in combination. At endpoint, tumor burden was calculated by counting peritoneal tumor nodules. **F)** The indicated CCNE1-high cells were treated with the IDH1 inhibitor GSK864 (IDH1i) and the DNA damaging agents olaparib (Olap) or cisplatin (Cis) alone (yellow) and in combination with cell permeable αKG (green) or citrate (blue). % of cells was assessed by crystal violet staining and normalized to controls. **G)** The indicated CCNE1-high cells were cultured under glutamine starvation conditions and treated with the DNA damaging agents olaparib (Olap) or cisplatin (Cis) alone (yellow) in combination with cell permeable αKG (green). % of cells was assessed by crystal violet staining and normalized to vehicle controls. **H)** Schematic of αKG-dependent dioxygenase CRISPR KO screen. **I)** Analysis of CRISPR KO screen. Shown is Log2 fold change of negative score in (CCNE1 + olaparib vs. CCNE1) vs. negative score in (EV + olaparib vs. EV). **J)** Venn diagram of the 5 negatively enriched genes in both CCNE1-high cell lines. **K)** The indicated CCNE1-high cells were transduced with shGFP (shContpurple) or two independent shRNAs targeting TMLHE (shTMLHE #1light blue, shTMLHE #2-dark blue) and treated with the DNA damaging agent olaparib (Olap) supplemented with cell permeable αKG or L-carnitine (L-Carn). % of cells was assessed by crystal violet staining and normalized to vehicle controls. **L)** The indicated CCNE1-high cells were treated with the carnitine synthesis inhibitor mildronate (Mildro) or the DNA damaging agent olaparib (Olap) alone (purple) and in combination (yellow). Combination treated cells were supplemented with cell permeable αKG (green) or L-carnitine (L-Carn; maroon). % of cells was assessed by crystal violet staining and normalized to vehicle controls. **M)** The indicated CCNE1-high cells were treated with the IDH1 inhibitor GSK864 (IDH1i) and the DNA damaging agent olaparib (Olap) alone (purple) and in combination (yellow). Combination treated cells were supplemented with L-carnitine (L-Carn; maroon). % of cells was assessed by crystal violet staining and normalized to vehicle controls. **N)** The indicated CCNE1-high cells were cultured under glutamine starvation conditions (purple) and treated with the DNA damaging agent olaparib (Olap) alone (yellow) or supplemented with L-carnitine (L-Carn; maroon). % of cells was assessed by crystal violet staining and normalized to controls. **O)** Schematic of αKG being upstream of TMLHE and carnitine. **(A-D, F)** Shown are representative data from at least 3 independent experiments in each isogenic cell line pair. **(G, K-N)** Shown are representative data from 2 independent experiments in each isogenic cell line pair. All graphs represent mean ± SD. ****p<0.005, ns = not significant

### αKG promotes carnitine synthesis

Next, we aimed to determine the contribution of αKG to *de novo* carnitine synthesis (schematic in **Fig. 2A**). Towards this goal, we performed mass spectrometry on cells that were depleted of αKG by glutamine starvation or treatment with the IDH1 inhibitor (**Fig. 1A and S1B**). Depletion of αKG using these methods corresponded to decreased abundance of L-carnitine as well as acetyl-carnitine and propionyl-carnitine, that was rescued by supplementation with αKG (**Fig. 2B-E and S2A-B**). We also observed a positive correlation between αKG and L-carnitine/acetyl-carnitine using cell line data from DepMap (**Fig. S2C**). These data provide evidence that cellular carnitine abundance can be regulated by αKG-dependent synthesis.

**Figure 2.**
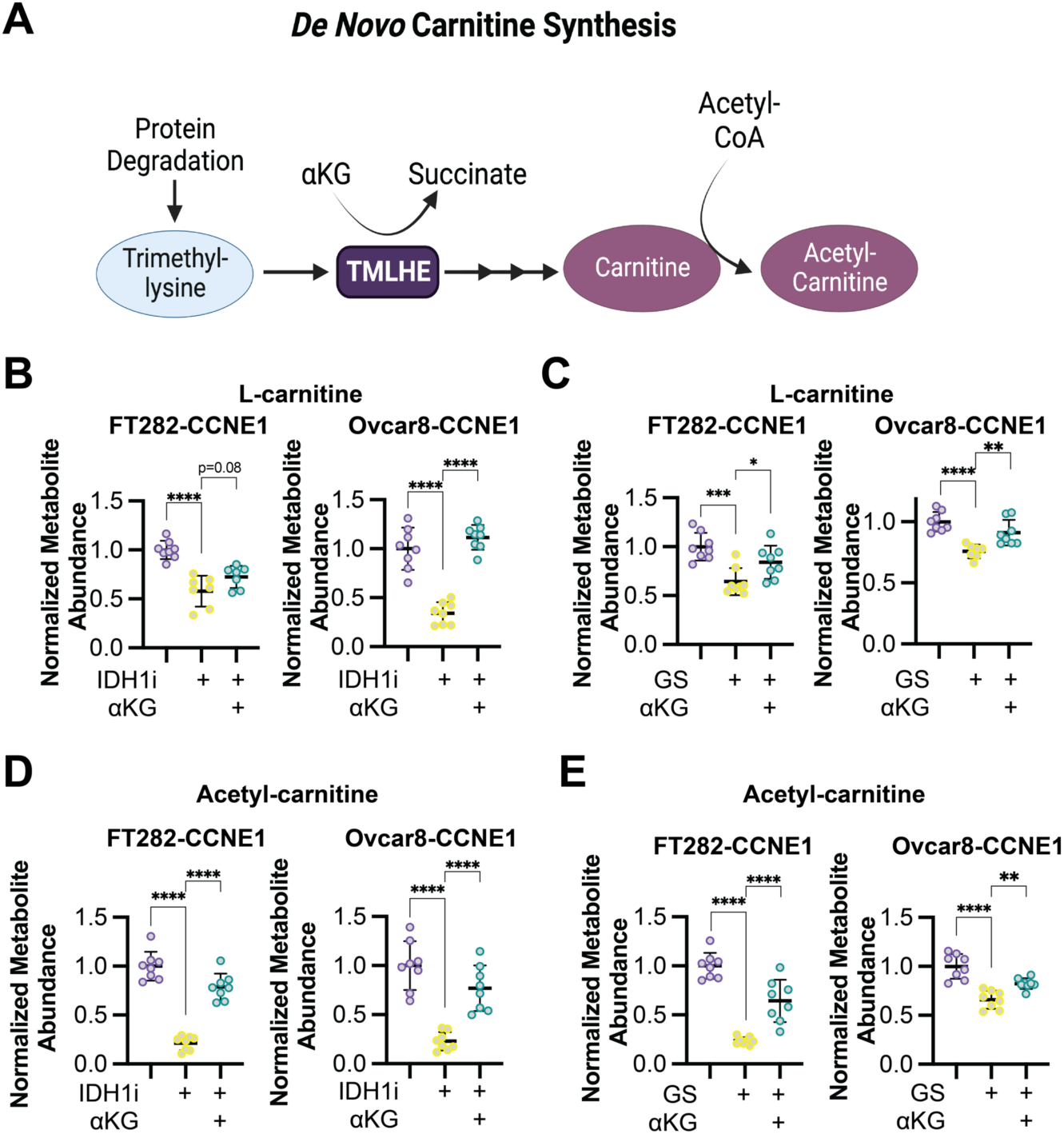
αKG promotes L-carnitine and acetyl-carnitine synthesis. **A)** Schematic of the *de novo* carnitine synthesis pathway. **B)** The indicated CCNE1-high cells were treated with the IDH1 inhibitor GSK864 (IDH1i) supplemented with cell permeable αKG, and L-carnitine abundance was assessed by LC-MS. Control = purple; IDH1i = yellow; αKG supplementation = green. **C)** The indicated CCNE1-high cells were cultured in normal media or under glutamine starvation conditions (GS) and supplemented with cell permeable αKG, and L-carnitine abundance was assessed by LC-MS. Control = purple; GS = yellow; αKG supplementation = green. **D)** Same as (A), but acetyl-carnitine was assessed by LC-MS. **E)** Same as (C), but acetyl-carnitine was assessed by LC-MS. Shown are representative data from 2-3 independent experiments in each cell line. All graphs represent mean ± SD. **p<0.01, ****p<0.001

### αKG-dependent carnitine synthesis via TMLHE increases histone acetylation

Carnitine is needed to transport some acyl-groups across the inner mitochondrial membrane, and acylcarnitines are known to be important for both energy production and redox balance (Flanagan et al., 2010; Longo et al., 2016). Surprisingly, only L-carnitine and acetylcarnitine but neither propionyl-carnitine nor butyryl-carnitine rescued the proliferation defect in response to DNA damage agents in combination with IDH1 inhibition, glutamine starvation, TMLHE knockdown, carnitine synthesis inhibition, or succinate supplementation (**Fig. S3A-E**). Similarly, the antioxidant n-acetyl-l-cysteine (NAC) also did not rescue the observed effects (**Fig. S3F-J**). Since these did not phenocopy the L-carnitine supplementation experiments (**Fig. 1K-N and S1**), these data indicate that energy production and/or ROS are not contributing to the observed effect. Recent publications also demonstrated that acetyl-carnitine is a precursor for nuclear acetyl-CoA that supports histone acetylation (Izzo et al., 2023; Kuna et al., 2023). Using a global mass spectrometry approach, we found that knockdown or inhibition of IDH1 decreased global histone acetylation to a greater extent than histone methylation (**Fig. 3A**). These data point to a previously unrecognized effect of αKG epigenetic rewiring via acetylation. This was confirmed in an additional cell line and with depletion of αKG or suppression of αKGDDs using knockdown/inhibition of IDH1, glutamine starvation, or succinate supplementation (**Fig. 3B-D and S4A)**. Total histone H3 and H4 acetylation was rescued by αKG supplementation (**Fig. 3B-D and S4A**). Decreased carnitine via knockdown of TMLHE also suppressed total histone H3 and H4 acetylation, which was not rescued by αKG (**Fig. 3E**).

**Figure 3.**
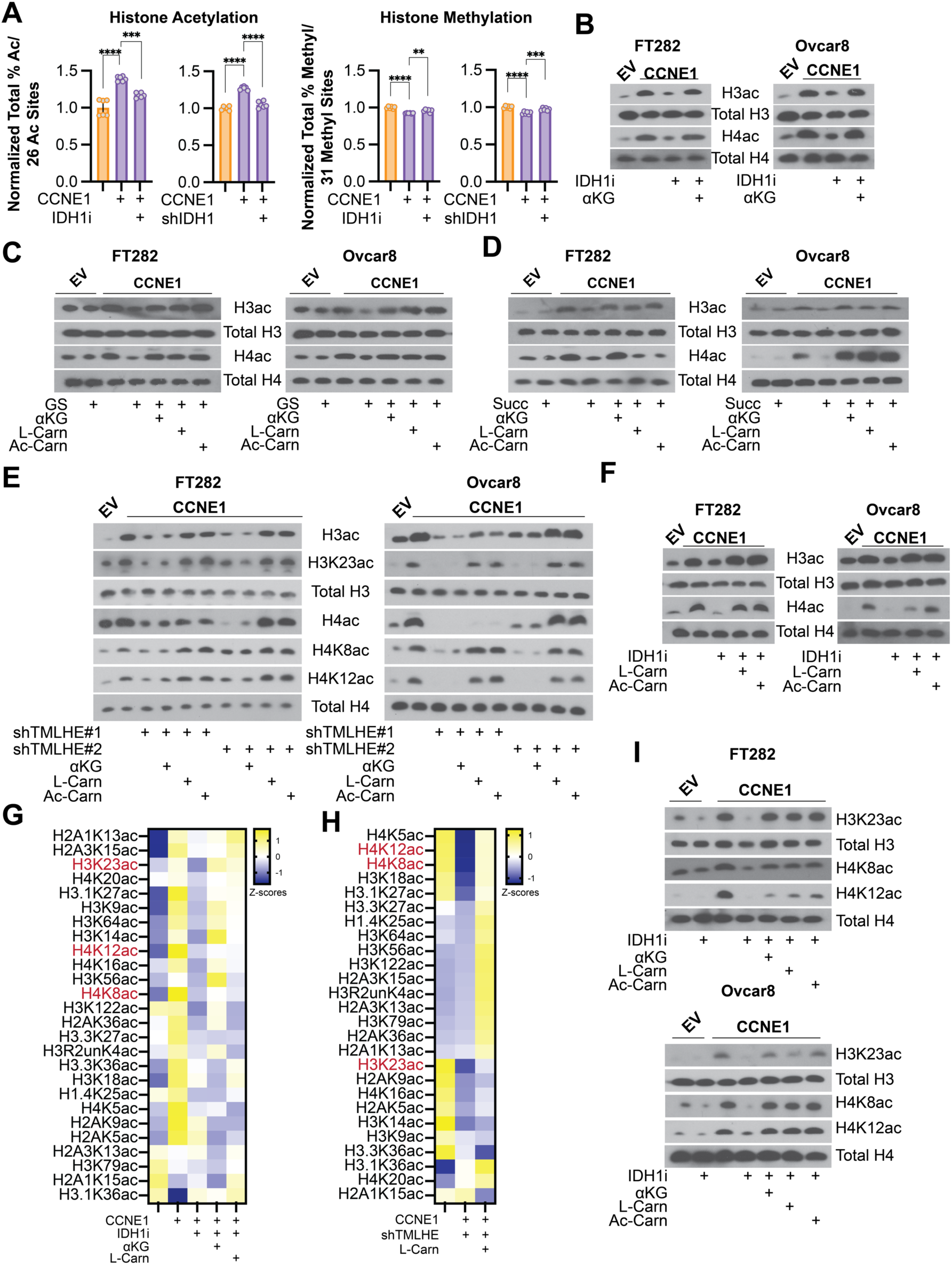
αKG-dependent *de novo* carnitine synthesis via TMLHE increases histone acetylation. **A)** Histone PTM abundance by mass spectrometry. Left-Sum of all % histone acetylation divided by total number of histone acetylation marks (26) in the indicated cells normalized to control. Right-Sum of all % histone methylation divided by total number of histone methylation marks (31) in the indicated cells normalized to control. Empty vector = orange, CCNE1-overexpressiong cells = purple. Graphs represent mean ± SD **p<0.01; ***p<0.005, ****p<0.001 **B)** The indicated cells were treated with the IDH1 inhibitor GSK864 (IDH1i) alone or supplemented with cell permeable αKG, and immunoblot analysis of the indicated proteins was performed. **C)** The indicated cells were cultured in normal media or under glutamine starvation conditions (GS) alone or supplemented with cell permeable αKG, L-carnitine (L-Carn), or O-acetyl-l-carnitine (Ac-Carn), and immunoblot analysis of the indicated proteins was performed. **D)** The indicated cells were supplemented with succinate with or without cell permeable αKG, L-carnitine (L-Carn), or O-acetyl-l-carnitine (Ac-Carn), and immunoblot analysis of the indicated proteins was performed. **E)** The indicated cells were transduced with two independent shRNAs targeting TMLHE (shTMLHE#1 and shTMLHE#2) and or supplemented with cell permeable αKG, L-carnitine (L-Carn), or O-acetyl-l-carnitine (Ac-Carn), and immunoblot analysis of the indicated proteins was performed. **F)** The indicated cells were treated with the IDH1 inhibitor GSK864 (IDH1i) alone or supplemented with cell permeable αKG, L-carnitine (L-Carn), or o-acetyl-l-carnitine (Ac-Carn), and immunoblot analysis of the indicated proteins was performed. **G-H)** Heatmap of histone acetylation marks in the indicated FT282 cells. Shown in red are marks that are significantly upregulated by CCNE1, downregulated by IDH1i or shTMLHE, and upregulated upon supplementation with αKG or L-carnitine. **I)** The indicated cells were treated with the IDH1 inhibitor GSK864 (IDH1i) alone or supplemented with cell permeable αKG, L-carnitine (L-Carn), or o-acetyl-l-carnitine (Ac-Carn), and immunoblot analysis of the indicated proteins was performed. Immunoblots shown are representative data from at least 2 independent experiments in each isogenic cell line pair.

We found that histone acetylation was rescued by either carnitine or acetyl-carnitine but only weakly by propionyl-carnitine or butyryl-carnitine (**Fig. 3C-F and S4B-D**), indicating that this rescue was specific to acetylation. Acetyl-carnitine can more directly contribute to acetylation reactions, whereas propionyl-carnitine or butyryl-carnitine contribute to propionylation and butyrylation, respectively (Trefely et al., 2020). Consistently, only L-carnitine and acetyl-carnitine rescued the decrease in proliferation in CCNE1-driven cells with suppressed αKG in combination with DNA damaging agents (**Fig. 1L-N, S1K-N, and S3A-D**). To gain a more global understanding of the specific histone acetylation marks that are regulated by αKG-mediated carnitine synthesis, we performed histone epiproteomics by mass spectrometry. Multiple acetylation marks were downregulated by IDH1 inhibition or knockdown (**Fig. 3G and S4E**). Using the IDH1 inhibitor, we further determine that multiple acetylation marks were rescued by αKG or L-carnitine (**Fig. 3G**). Similarly, knockdown of TMLHE decreased multiple histone acetylation marks, many of which were rescued by L-carnitine (**Fig. 3H**). Notably, addition of carnitine or knockdown of TMLHE did not have marked effects on histone methylation (**Fig. S4F**), indicating methylation reactions are dispensable for the observed phenotype. Cross-comparison of these datasets identified 3 specific histone acetyl marks – H3K23ac, H4K8ac and H4K12ac – that were specifically sensitive to regulation by this axis and validated by western blotting in multiple cell lines under different conditions (**Fig. 3E and 3I**). Together, our data provide evidence that αKG contributes to histone acetylation by acting as a co-substrate for the carnitine synthesis enzyme TMLHE.

### αKG-mediated carnitine production and histone acetylation suppresses DNA damage via HR

CCNE1-driven cells are known to accumulate DNA damage and require proficient HR to survive (Etemadmoghadam et al., 2013). Indeed, we observed an increase in DNA damage foci in CCNE1 overexpressing cells similar to the positive control cisplatin (**Fig. S5A-B**). Knockdown or inhibition of IDH1 at timepoints and doses that do not markedly affect proliferation (**Fig. S5C-E**) increases DNA damage markers in CCNE1 overexpressing cells but not controls (**Fig. 4A and S5F**). This was rescued by αKG, L-carnitine, and acetyl-carnitine, but not citrate, propionyl-carnitine, or butyryl-carnitine (**Fig. 4A and S5G**). Consistent with the role of αKG in carnitine synthesis, we also observed an increase in DNA damage foci upon TMLHE knockdown, which was rescued by L-carnitine and acetylcarnitine, but not αKG, propionyl-carnitine, or butyryl-carnitine (**Fig. 4B and S4H**). Together, these data indicate the observed effects are due to an acetylation reaction. Prior studies have provided a link between histone acetylation and DNA repair proficiency (Gong and Miller, 2013; Song et al., 2023), and we observed an increase in histone acetylation downstream of αKG-mediated TMLHE (**Fig. 3**). Histone acetylation can affect DNA repair in multiple ways, including directly through promoting chromatin relaxation and recruitment of specific DNA repair factors, or indirectly through promoting transcription of DNA repair genes. Using a *Xenopus* egg extract model (Barrows et al., 2022), we determined that IDH1 inhibition increased DSB formation and decreased HR, which was rescued by αKG (**Fig. 4C-D**). This model uncouples DNA repair from gene expression, demonstrating this is likely a direct effect on chromatin, not changes in transcription of DNA repair factors. Using a model of induced DSBs in an endogenously CCNE1 overexpressing cell line (Tang et al., 2013), we found increased DSBs upon IDH1i treatment, which was rescued by αKG and L-carnitine (**Fig. 4E**). We also observed decreased BRCA1 recruitment to DSBs in cells treated with the IDH1i, which was rescued by αKG, L-carnitine, and acetyl-carnitine (**Fig. 4F**). Notably, this was not due to decreased S phase as these cells had similar BrdU incorporation regardless of the treatment (**Fig. S5I**). We identified H4K8, H3K23, and H4K12 histone acetylation marks to be specifically regulated by αKG-mediated carnitine synthesis (**Fig. 3**). ChIP demonstrated decreased occupancy of all three marks at DSBs in IDH1i-treated cells in addition to increased γH2AX (**Fig. 4G and S5J**). The most robust effects were with H4K8ac, which was rescued by both αKG and L-carnitine (**Fig. 4G**). Finally, we found that TMLHE knockdown phenocopied IDH1 inhibition by increasing γH2AX and decreasing H4K8ac occupancy at DSBs (**Fig. 4H**). This was only rescued by carnitine, not by αKG. Our data provide evidence that IDH1-mediated αKG production is required for HR in CCNE1-driven models, and this is through TMLHE and carnitine-mediated histone acetylation (**Fig. 4I**).

**Figure 4.**
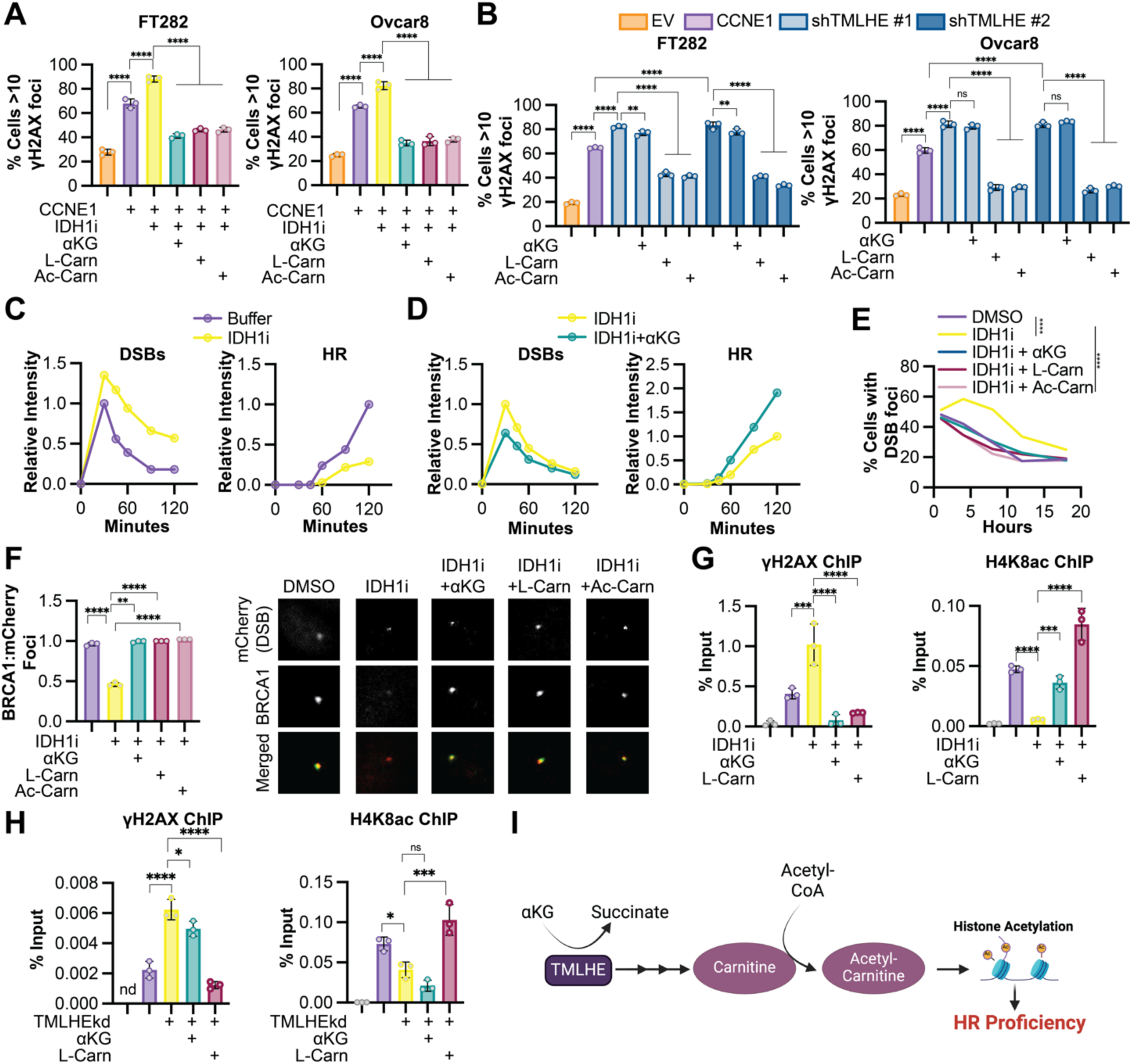
αKG-mediated carnitine production is required for DNA double strand break repair in CCNE1-driven models. **A)** The indicated cells (EV = orange, CCNE1 = purple) were treated with the IDH1 inhibitor GSK864 (IDH1i) alone (yellow) or supplemented with cell permeable αKG (green), L-carnitine (L-Carn, maroon) or o-acetyl-l-carnitine (Ac-Carn, light maroon), and immunofluorescence analysis for γH2AX foci was performed. **B)** The indicated cells (EV = orange, CCNE1 = purple) were transduced with two independent shRNAs targeting TMLHE (shTMLHE#1, light blue; and shTMLHE#2, dark blue) and or supplemented with cell permeable αKG, L-carnitine (L-Carn), or o-acetyl-l-carnitine (Ac-Carn), and immunofluorescence analysis for γH2AX foci was performed. **C-D)** *Xenopus* egg extract experiments. Plasmid DNA was replicated in *Xenopus* egg extract −/+ IDH1i. After 45 min, reactions were supplemented with AgeI (+0 min) to generate a DSB. Samples were withdrawn at the indicated times, resolved by agarose gel electrophoresis, and band intensity was quantified. **C)** Quantification of linear (DSBs) and high molecular weight (HR) intermediates in control (purple) and IDH1i treated (yellow) extracts. **D)** Quantification of linear (DSBs) and high molecular weight (HR) intermediates in control (purple) and IDH1i treated (yellow) extracts. **E)** U2OS-mCherry-LacI-Fok1 cells were treated with the IDH1 inhibitor GSK864 (IDH1i) alone (yellow) or supplemented with cell permeable αKG (green), L-carnitine (L-Carn, maroon), or o-acetyl-l-carnitine (Ac-Carn, light maroon), and number of mCherry DSB foci was counted over time. **F)** Same as (E) but co-localized of BRCA1 foci and mCherry DSB foci were quantified. Representative images of colocalization of mCherry DSB foci (red) and BRCA1 foci (green). **G)** Same as (E) but chromatin immunoprecipitation (ChIP) was performed with γH2AX or H4K8ac antibodies. **H)** U2OS-mCherry-LacI-Fok1 cells were transduced with shRNA targeting TMLHE (shTMLHE) alone and supplemented with cell permeable αKG or L-carnitine (L-Carn), and chromatin immunoprecipitation (ChIP) was performed with γH2AX or H4K8ac antibodies. **I)** Schematic of this study illustrating αKG-mediated *de novo* carnitine synthesis via TMLHE leading to HR proficiency via histone acetylation. Shown are representative data from 3 independent experiments in each cell line or egg extract. Graphs represent mean ± SD. **p<0.01; ***p<0.005; ns = not significant; nd = no data. ****p<0.001

## Discussion

αKG is well known for its role in DNA and histone demethylation, via action as a co-substrate for αKGDDs (Islam et al., 2018). Here, we report that αKG drives acetylation to promote HR-mediated DNA DSB repair. We discovered that αKG-dependent histone acetylation occurs via TMLHE-mediated *de novo* carnitine synthesis as TMLHE is a αKGDD. This study provides a new link between αKG and histone acetylation and opens up new avenues of exploration into the epigenetic and translational consequences of altered αKG levels.

Epigenetic modifications such as histone PTMs are metabolically sensitive (Izzo et al., 2021). Prior work has shown that depletion of αKG, for instance via mutations in IDH1/2 that instead produce the oncometabolite D2-HG, inhibits αKGDD activity leading to changes in DNA and histone methylation (Inoue et al., 2016; Molenaar et al., 2018; Sulkowski et al., 2017; Sulkowski et al., 2020; Sulkowski et al., 2018; Wang et al., 2015). To our knowledge, this is the first report of αKG promoting histone acetylation. Histone acetylation is partly controlled by nuclear-cytoplasmic acetyl-CoA pools (Sivanand et al., 2017), and recent work has demonstrated that the endogenous acyl-carrier L-carnitine plays an active role in histone acetylation by shuttling acetyl units in the form of acetylcarnitine from the mitochondria and/or peroxisome and acting as a nuclear acetyl-CoA precursor (Izzo et al., 2023; Kuna et al., 2023). In this study, we have now identified an upstream regulator of the carnitine-histone acetylation axis via the αKGDD TMLHE. We posit that acetylation events are likely affected in these other biological scenarios such as IDH1/2 mutations where αKG is depleted.

Studies have found that inhibiting αKGDD activity due to mutations in IDH1/2, FH, or SDH increases sensitivity to DNA damaging agents, and the mechanisms described are due to changes in methylation (Inoue et al., 2016; Jiang et al., 2015; Molenaar et al., 2018; Sulkowski et al., 2017; Sulkowski et al., 2020; Sulkowski et al., 2018; Wang et al., 2015). Consistently, we found that depleting αKG either through inhibition of wildtype IDH1 or glutamine starvation, or inhibiting αKGDD activity via exogenous succinate, sensitized CCNE1-driven cells to DNA damaging agents (**Fig. 1**). What our study shows for the first time is that this phenomenon is via histone acetylation and subsequent DNA repair, whereas histone methylation seems to be dispensable (**Fig. 3-4**). Indeed, histone acetylation is implicated in promoting DNA damage response in multiple ways (Song et al., 2023). Histone acetylation plays a major role in regulating the chromatin structure around the DSB that is required for both chromatin relaxation and efficient recruitment of DSB repair proteins as well as promoting transcription of DNA repair genes. Our data using a *Xenopus* egg extract system suggested αKG-mediated HR was a more direct effect of histone acetylation on either chromatin relaxation or recruitment of repair factors. Interestingly, we observed multiple acetylation sites that were sensitive to αKG-mediated carnitine synthesis, including H4K8ac, H4K12ac, and H3K23ac (**Fig. 3**). Prior work has found that acetylation of the H4 tail is important for direct HR-mediated repair (Aricthota et al., 2022; Gong and Miller, 2013), and we observed increased HR by increased BRCA1 co-localization with DSBs in cells with high αKG and carnitine (**Fig. 4F**). Less is understood about H3K23ac in DNA repair. It is possible that these acetylation marks also help to promote chromatin relaxation or other DNA damage pathways, which will be investigated in future studies. Notably, low H4K12ac was previously shown to be associated with HR-deficiency in ovarian cancer patients (McDermott et al., 2020), further illustrating the translational potential of our work.

CCNE1 is an oncogene that is amplified or overexpressed in a variety of human cancers including HGSOC (Karst et al., 2014; Kuhn et al., 2016). These cancers are typically resistant to standard-of-care DNA damaging agents and alternative therapies are limited (Etemadmoghadam et al., 2009; Patch et al., 2015). Our data suggest that inhibition of wildtype IDH1, depletion of αKG through glutamine restriction, or inhibition of carnitine synthesis may act as a novel therapeutic approach in these cancers. There is evidence in other cancer types that point towards using wildtype IDH1 inhibition in combination with chemotherapy as a strategy for treatment, although these studies focused on redox balance and ROS (Vaziri-Gohar et al., 2022; Zarei et al., 2022; Zarei et al., 2023). Glutamine starvation (or glutaminase inhibition) has also been shown to have similar effects, due in part to decreased GSH and redox imbalance (Gross et al., 2014; Guo et al., 2021; Mukhopadhyay et al., 2020). We did not see rescue using the antioxidant NAC, suggesting multiple context-dependent mechanisms are at play. Carnitine is also an essential part of the diet and is enriched in foods such as milk, meat, fish, and cheese (Steiber et al., 2004). It will therefore be interesting in future studies to determine the contribution of dietary carnitine to histone acetylation and HR-proficiency and whether modulating dietary carnitine would affect response of tumors to DNA damaging agents or even slow the development of tumors.

In summary, we found that αKG was required for histone acetylation via TMHLE-catalyzed carnitine synthesis to facilitate HR-mediated DNA repair in CCNE1-driven models. We speculate that this provides targetable insight into the metabolic mechanism of HR-proficient resistance to DNA damaging agents. Moreover, our data provide evidence of an unappreciated pathway for αKG to promote carnitine synthesis that supports histone acetylation in addition to its known roles in demethylation. This should prompt re-appraisal of the role of αKG in physiology and disease via mechanisms other than the αKG-dependent DNA and histone demethylases.

## Acknowledgements

We would like to thank Dr. Kathryn Wellen (University of Pennsylvania) for critical reading of our manuscript, Fran Vasquez for help with the CRISPR schematic, Uma Chandran and Jiefei Wang (UPMC Hillman Cancer Center Bioinformatics Core) for help with bioinformatic analysis of the CRISPR screen, and Maureen Lyons (UPMC Hillman Cancer Center Cancer Genomics Core) for help with sequencing of the CRISPR library. Multiple schematics created with BioRender.com. This work was supported by National Institutes of Health (R37CA240625 to KMA, R01CA259111 to KMA and NWS, T32GM133332 to ARC, R01CA242021 to NH, R35GM119512 to DTL), the American Cancer Society (RSG-19-113-01-CCG to KMA), the Ovarian Cancer Research Alliance (MIG-2023-2-1018 to AU), HERA Ovarian Cancer Foundation (AU), the Melanoma Research Foundation (RB), the Hollings Cancer Center Abney Graduate Fellowship (JAD), and the UPMC Hillman Cancer Center. This project used the Hillman Animal Facility, Cancer Genomics Facility, and the Cancer Bioinformatics Services that are supported in part by award P30CA047904.

## Author Contributions

Apoorva Uboveja: Conceptualization, Investigation, Methodology, Writing – Original Draft, Visualization, Writing – Review & Editing, Funding Acquisition. Zhentai Huang: Investigation, Methodology. Raquel Buj: Investigation, Methodology, Writing – Review & Editing. Amandine Amalric: Investigation, Methodology, Writing – Review & Editing. Hui Wang: Investigation. Naveen Kumar Tangudu: Investigation, Writing – Review & Editing. Aidan R. Cole: Investigation, Writing – Review & Editing. Emily Megill: Investigation. Daniel Kantner: Investigation. Adam Chatoff: Investigation. Hafsah Ahmad: Investigation. Mariola M. Marcinkiewicz: Investigation. Julie A. Disharoon: Investigation. Sarah Graff: Investigation. Erika S. Dahl: Investigation. Nadine Hempel: Writing – Review & Editing. Wayne Stallaert: Investigation, Methodology, Writing – Review & Editing. Simone Sidoli: Investigation, Methodology, Writing – Review & Editing, Supervision. Benjamin G. Bitler: Investigation, Methodology, Writing – Review & Editing. David T. Long: Investigation, Methodology, Writing – Review & Editing, Supervision. Nathaniel W. Snyder: Conceptualization, Investigation, Methodology, Writing – Original Draft, Visualization, Writing – Review & Editing, Supervision, Funding Acquisition. Katherine M. Aird: Conceptualization, Visualization, Writing – Original Draft, Writing – Review & Editing, Supervision, Project Administration, Funding Acquisition.

## Declaration of Interests

All authors declare no competing interests.

## Supplemental Figures

**Figure S1.**
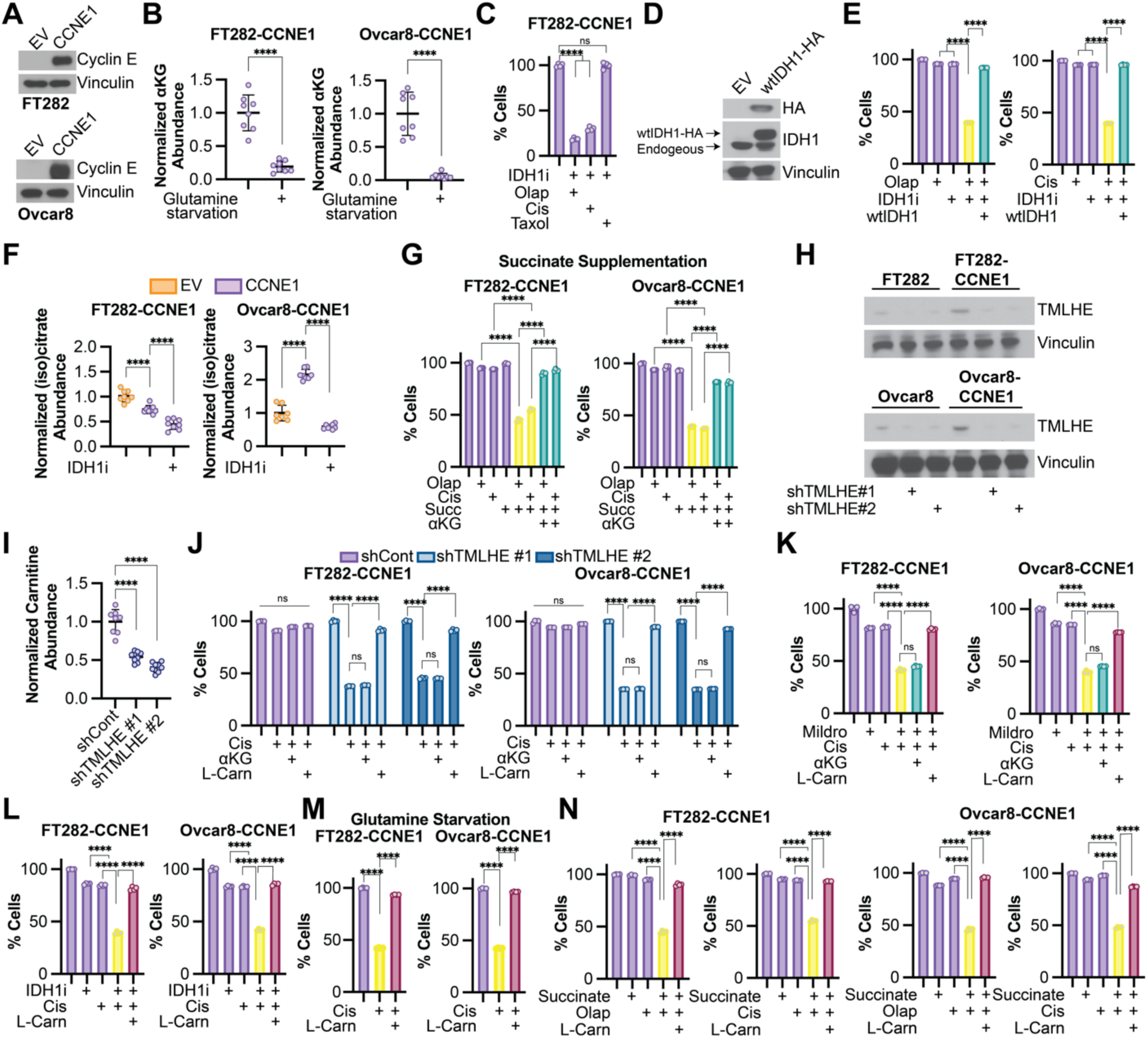
IDH1 inhibitor, succinate supplementation, TMLHE knockdown, and carnitine synthesis inhibition decrease proliferation in response to DNA damaging agents in CCNE1-driven models, which is rescued by carnitine supplementation. Related to Figure 1. **A)** Immunoblot analysis of cyclin E in the indicated isogenic cells. Vinculin was used as a loading control. **B)** The indicated cells were cultured in normal media or under glutamine starvation conditions, and αKG and abundance was assessed by LC-MS. **C)** FT282-CCNE1 cells were treated with the IDH1 inhibitor GSK864 (IDH1i) in combination with the DNA damaging agents olaparib (Olap) or cisplatin (Cis) or the microtubule stabilizer paclitaxel (Taxol). % of cells was assessed by crystal violet staining and normalized to controls. **D)** Immunoblot analysis of IDH1 and HA in wtIDH1-3XHA transduced Ovcar8 cells. Vinculin was used as a loading control. **E)** FT282-CCNE1 cells were transduced with empty vector or a wildtype IDH1 overexpression plasmid (wtIDH1). Cells were treated with the IDH1 inhibitor GSK864 (IDH1i) and DNA damaging agents olaparib (Olap) or cisplatin (Cis) alone (purple) or in combination (yellow). wtIDH1 overexpression rescued proliferation (green). % of cells was assessed by crystal violet staining and normalized to controls. **F)** The indicated cells were treated with the IDH1 inhibitor GSK864 (IDH1i), and (iso)citrate abundance was assessed by LC-MS. Cells expressing empty vector (EV) = orange; Cells expressing CCNE1 (CCNE1) = purple. **G)** The indicated CCNE1-high cells were treated with succinate and DNA damaging agents olaparib (Olap) and cisplatin (Cis) alone (purple) or in combination (yellow). Combination treated cells were supplemented with cell permeable αKG (green). % of cells was assessed by crystal violet staining and normalized to controls. **H)** The indicated cells were transduced with two independent shRNAs targeting TMLHE (shTMLHE #1 and shTMLHE #2). Immunoblot analysis of TMLHE. Vinculin was used as a loading control. **I)** The indicated cells were transduced with two independent shRNAs targeting TMLHE (shTMLHE #1 and shTMLHE #2), and carnitine was assessed by LC-MS. **J)** The indicated CCNE1-high cells were transduced with shGFP (shCont-purple) or two independent shRNAs targeting TMLHE (shTMLHE #1-light blue, shTMLHE #2-dark blue) and treated with the DNA damaging agent olaparib (Olap) supplemented with cell permeable αKG or L-carnitine (L-Carn). % of cells was assessed by crystal violet staining and normalized to controls. **K)** The indicated cells were treated with the carnitine synthesis inhibitor mildronate (Mildro), and the DNA damaging agent cisplatin (Cis) alone (purple) and in combination (yellow). Combination treated cells were supplemented with cell permeable αKG (green) or L-carnitine (L-Carn; maroon). % of cells was assessed by crystal violet staining and normalized to controls. **L)** The indicated CCNE1-high cells were treated with the IDH1 inhibitor GSK864 (IDH1i) and the DNA damaging agent cisplatin (Cis) alone (purple) and in combination (yellow). Combination treated cells were supplemented with L-carnitine (L-Carn; maroon). % of cells was assessed by crystal violet staining and normalized to controls. **M)** The indicated CCNE1-high cells were cultured under glutamine starvation conditions (purple) and treated with the DNA damaging agent cisplatin (Cis) alone (yellow) or supplemented with L-carnitine (L-Carn; maroon). % of cells was assessed by crystal violet staining and normalized to controls. **N)** The indicated CCNE1-high cells were treated with succinate and DNA damaging agents olaparib (Olap) and cisplatin (Cis) alone (purple) or in combination (yellow). Combination treated cells were supplemented with L-carnitine (L-Carn; maroon). % of cells was assessed by crystal violet staining and normalized to controls. **(A, D-F)** Shown are representative data from at least 3 independent experiments in each isogenic cell line pair. **(B-C, G-N)** Shown are representative data from 2 independent experiments in each isogenic cell line pair. All graphs represent mean ± SD. **p<0.01, ****p<0.001, ns = not significant

**Figure S2.**
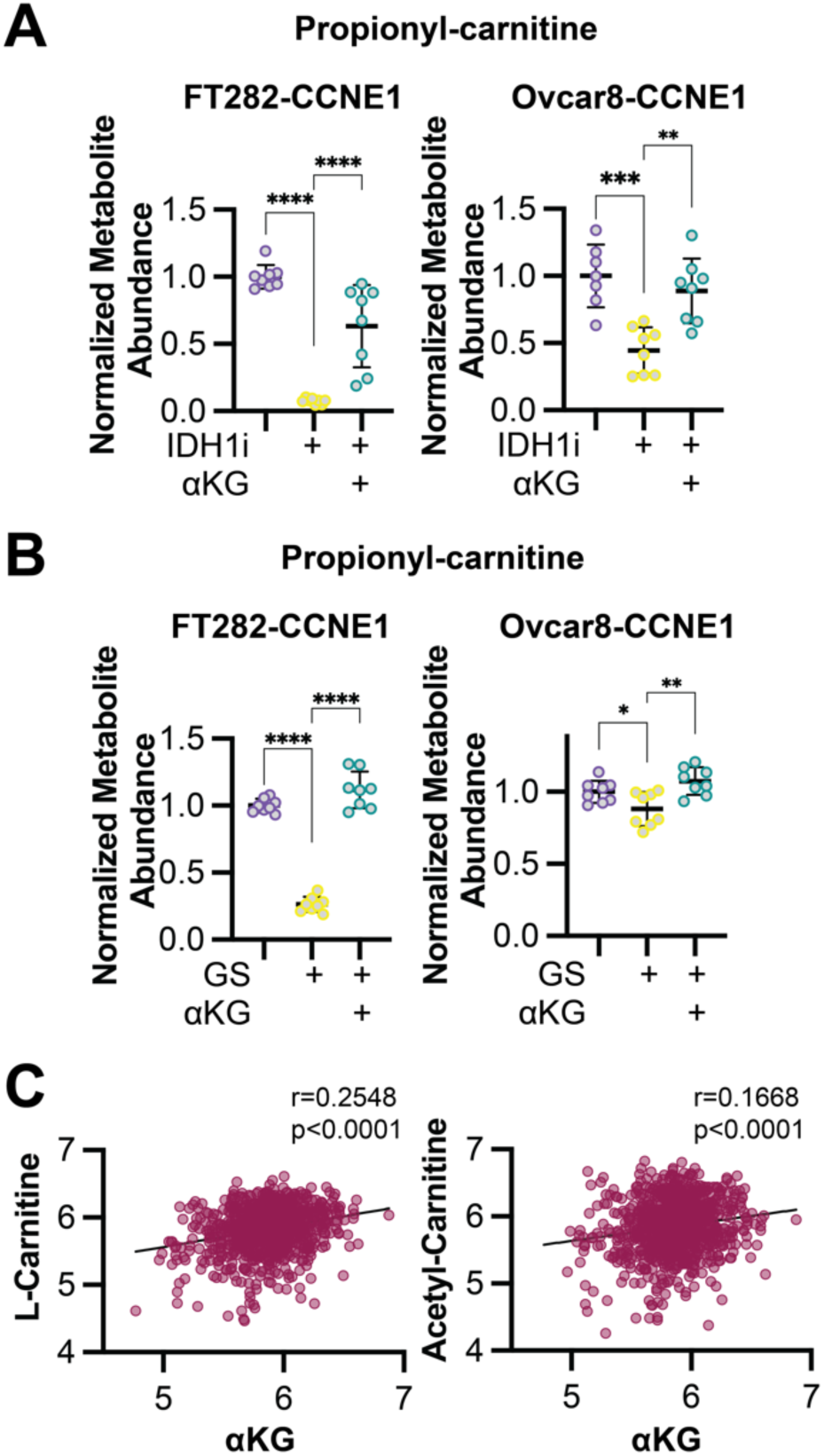
αKG promotes propionyl-carnitine synthesis, and publicly available data show correlation between αKG and carnitine/acetyl-carnitine. Related to Figure 2. **A)** The indicated CCNE1-high cells were treated with the IDH1 inhibitor GSK864 (IDH1i) supplemented with cell permeable αKG, and propionyl-carnitine abundance was assessed by LC-MS. Control = purple; IDH1i = yellow; αKG supplementation = green. **B)** The indicated CCNE1-high cells were cultured in normal media or under glutamine starvation conditions (GS) and supplemented with cell permeable αKG, and propionyl-carnitine abundance was assessed by LC-MS. Control = purple; GS = yellow; αKG supplementation = green. Shown are representative data from 2-3 independent experiments in each cell line. All graphs represent mean ± SD. **p<0.01, ****p<0.001 **C)** Metabolite abundance data in all cell lines taken from DepMap.

**Figure S3.**
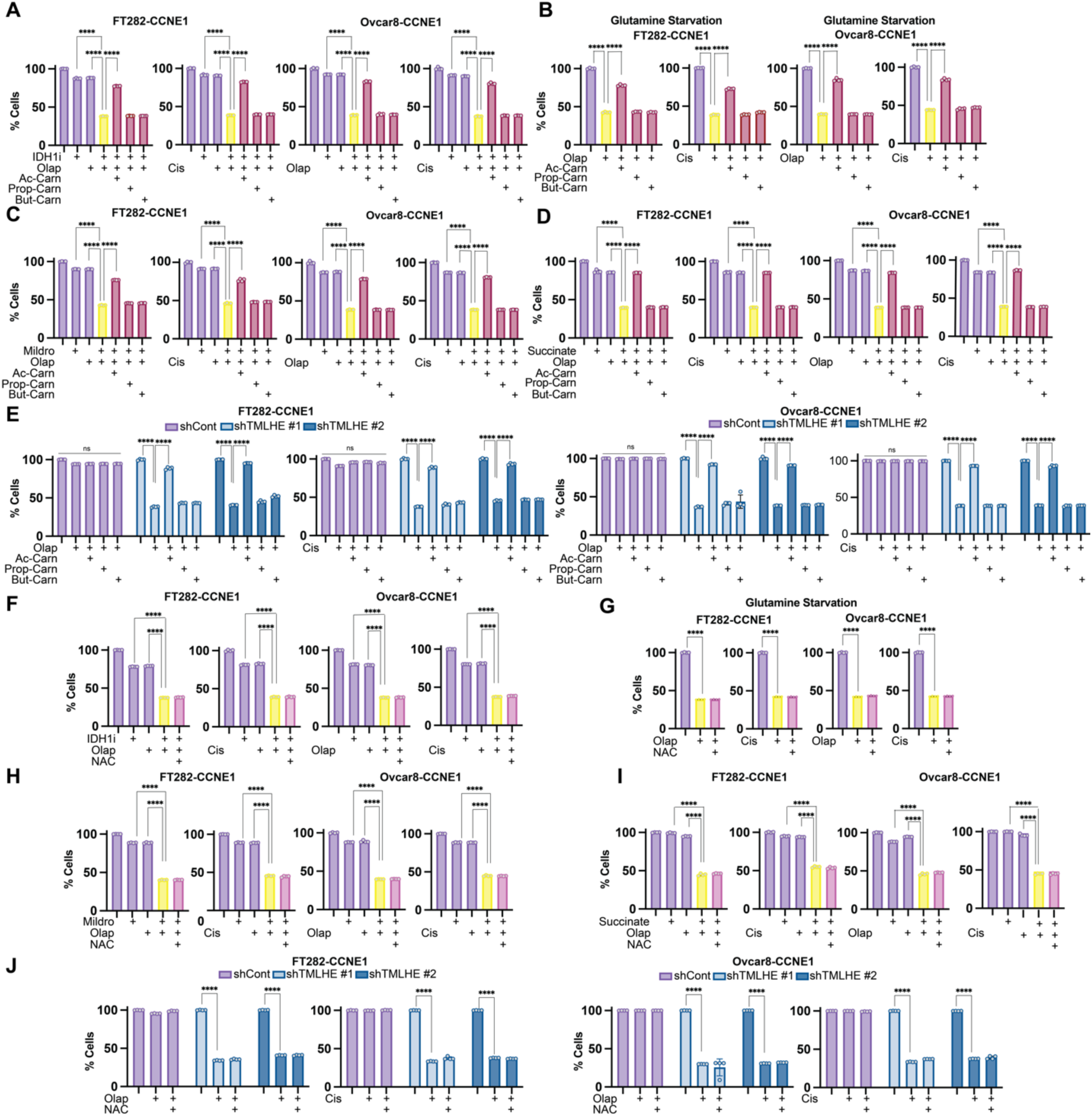
Butyryl- and propionyl-carnitine or n-acetyl-l-cysteine do not rescue decreased αKG-mediated proliferation defects in combination with DNA damaging agents, TMLHE knockdown, or carnitine synthesis inhibition. Related to Figure 3. For all experiments, % of cells was assessed by crystal violet staining and normalized to controls. **A)** The indicated cells CCNE1 overexpressing cells were treated with IDH1 inhibitor GSK864 (IDH1i) and DNA damaging agents olaparib (Olap) or cisplatin (Cis) alone (purple) and in combination (yellow) and supplemented with o-acetyl-l-carnitine (Ac-Carn, maroon), propionyl-carnitine (Prop-Carn, maroon), or butyryl-carnitine (But-Carn, maroon). **B)** The indicated CCNE1 overexpressing cells were cultured under glutamine starvation conditions and treated with the DNA damaging agents olaparib (Olap) or cisplatin (Cis) alone (yellow) or supplemented with o-acetyl-l-carnitine (Ac-Carn, maroon), propionyl-carnitine (Prop-Carn, maroon), or butyryl-carnitine (But-Carn, maroon). **C)** The indicated CCNE1 overexpressing cells were treated with the carnitine synthesis inhibitor mildronate (Mildro) and the DNA damaging agents olaparib (Olap) or cisplatin (Cis) alone (purple) and in combination (yellow) and supplemented with o-acetyl-l-carnitine (Ac-Carn, maroon), propionyl-carnitine (Prop-Carn, maroon), or butyryl-carnitine (But-Carn, maroon). **D)** The indicated CCNE1 overexpressing cells were supplemented with succinate and treated with the DNA damaging agents olaparib (Olap) or cisplatin (Cis) alone (purple) and in combination (yellow) and supplemented with o-acetyl-l-carnitine (Ac-Carn, maroon), propionyl-carnitine (Prop-Carn, maroon), or butyryl-carnitine (But-Carn, maroon). **E)** The indicated CCNE1 overexpressing cells were transduced with two independent shRNAs targeting TMLHE (shTMLHE#1-light blue, shTMLHE#2-dark blue) and treated with the DNA damaging agents olaparib (Olap) or cisplatin (Cis) alone or supplemented with o-acetyl-l-carnitine (Ac-Carn), propionyl-carnitine (Prop-Carn), or butyryl-carnitine (But-Carn). **F)** The indicated CCNE1 overexpressing cells were treated with IDH1 inhibitor GSK864 (IDH1i) and DNA damaging agents olaparib (Olap) or cisplatin (Cis) alone (purple) and in combination (yellow) and supplemented with n-acetyl-l-cysteine (NAC, pink). **G)** The indicated CCNE1 overexpressing cells were cultured in normal media or under glutamine starvation conditions and treated with the DNA damaging agents olaparib (Olap) or cisplatin (Cis) alone (yellow) and supplemented with n-acetyl-l-cysteine (NAC, pink). **H)** The indicated CCNE1 over-expressing cells were treated with the carnitine synthesis inhibitor mildronate (Mildro) and the DNA damaging agents olaparib (Olap) or cisplatin (Cis) alone (purple) and in combination (yellow) and supplemented with n-acetyl-l-cysteine (NAC, pink). **I)** The indicated CCNE1 overexpressing cells were supplemented with succinate and treated with the DNA damaging agents olaparib (Olap) or cisplatin (Cis) alone (purple) and in combination (yellow) and supplemented with nacetyl-l-cysteine (NAC, pink). **J)** The indicated cells CCNE1 overexpressing cells were transduced with two independent shRNAs targeting TMLHE (shTMLHE#1-light blue, shTMLHE#2-dark blue) and treated with the DNA damaging agents olaparib (Olap) or cisplatin (Cis) alone or supplemented with n-acetyl-l-cysteine (NAC). Shown are representative data from 2-3 independent experiments in each cell line. All graphs represent mean ± SD. ****p<0.001, ns = not significant

**Figure S4.**
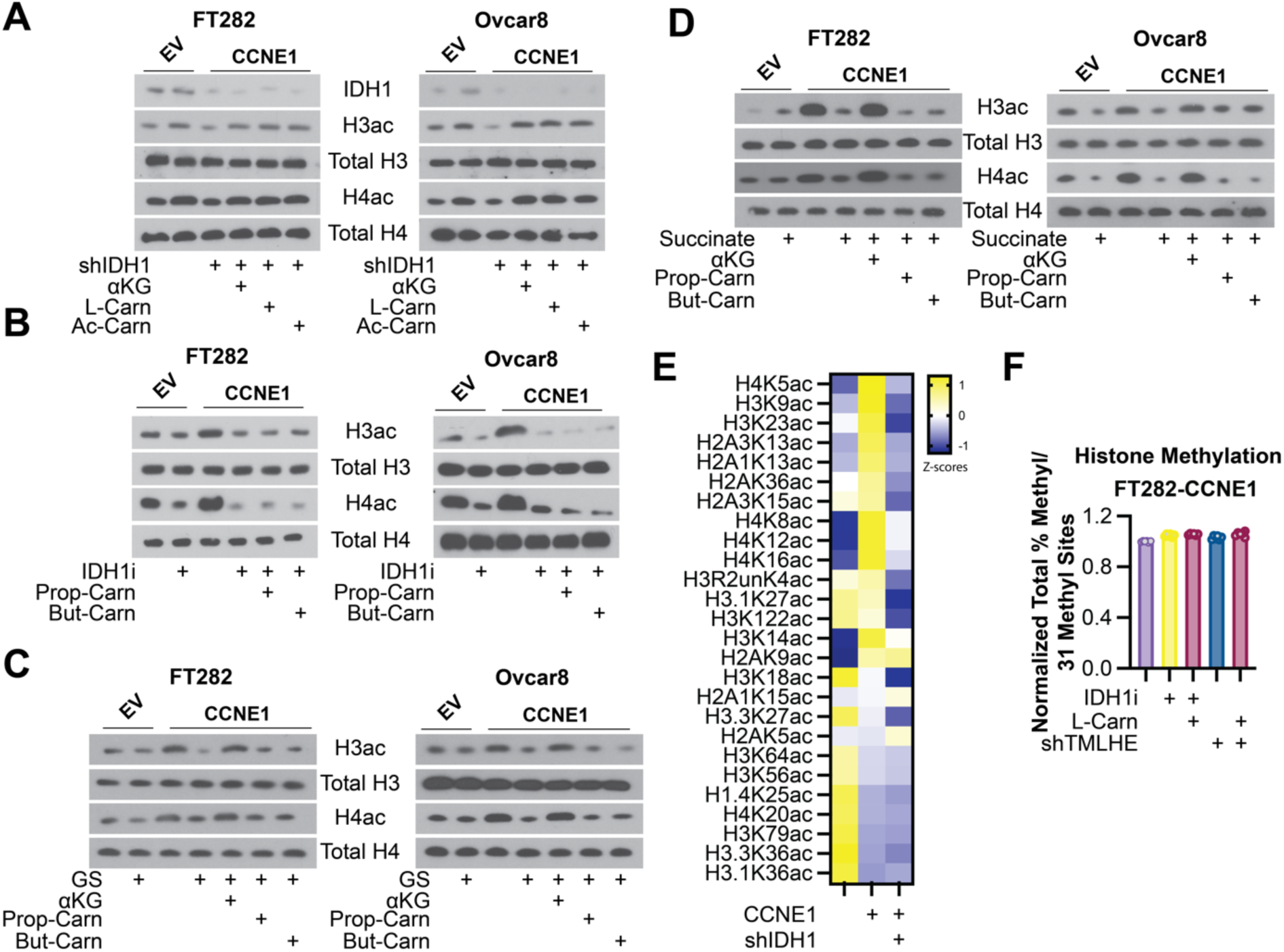
Histone acetylation is not rescued by propionyl-carnitine or butyryl-carnitine, and αKG production affects histone acetylation more robustly than histone methylation. Related to Figure 3. **A)** The indicated cells were transduced with shRNA targeting IDH1 and supplemented with cell permeable αKG, L-carnitine, or O-acetyl carnitine, and immunoblot analysis of the indicated proteins was performed. **B)** The indicated cells were treated with IDH1 inhibitor GSK864 (IDH1i) and supplemented with propionyl-carnitine (Prop-Carn) or butyryl-carnitine (But-Carn), and the indicated proteins were assessed by immunoblotting. **C)** The indicated cells were cultured in normal media or under glutamine starvation conditions (GS) and supplemented with cell permeable αKG, propionyl-carnitine (Prop-Carn), or butyryl-carnitine (But-Carn), and the indicated proteins were assessed by immunoblotting. **D)** The indicated cells were supplemented with succinate and cell permeable αKG, propionyl-carnitine (Prop-Carn), or butyryl-carnitine (But-Carn), and the indicated proteins were assessed by immunoblotting. **E)** Heatmap of histone acetylation marks in FT282 cells upon IDH1 knockdown. **F)** Sum of all % histone methylation divided by total number of histone methylation marks (31) in the indicated cells normalized to control. Graphs represent mean ± SD. Shown are representative data from 2-3 independent experiments in each cell line.

**Figure S5.**
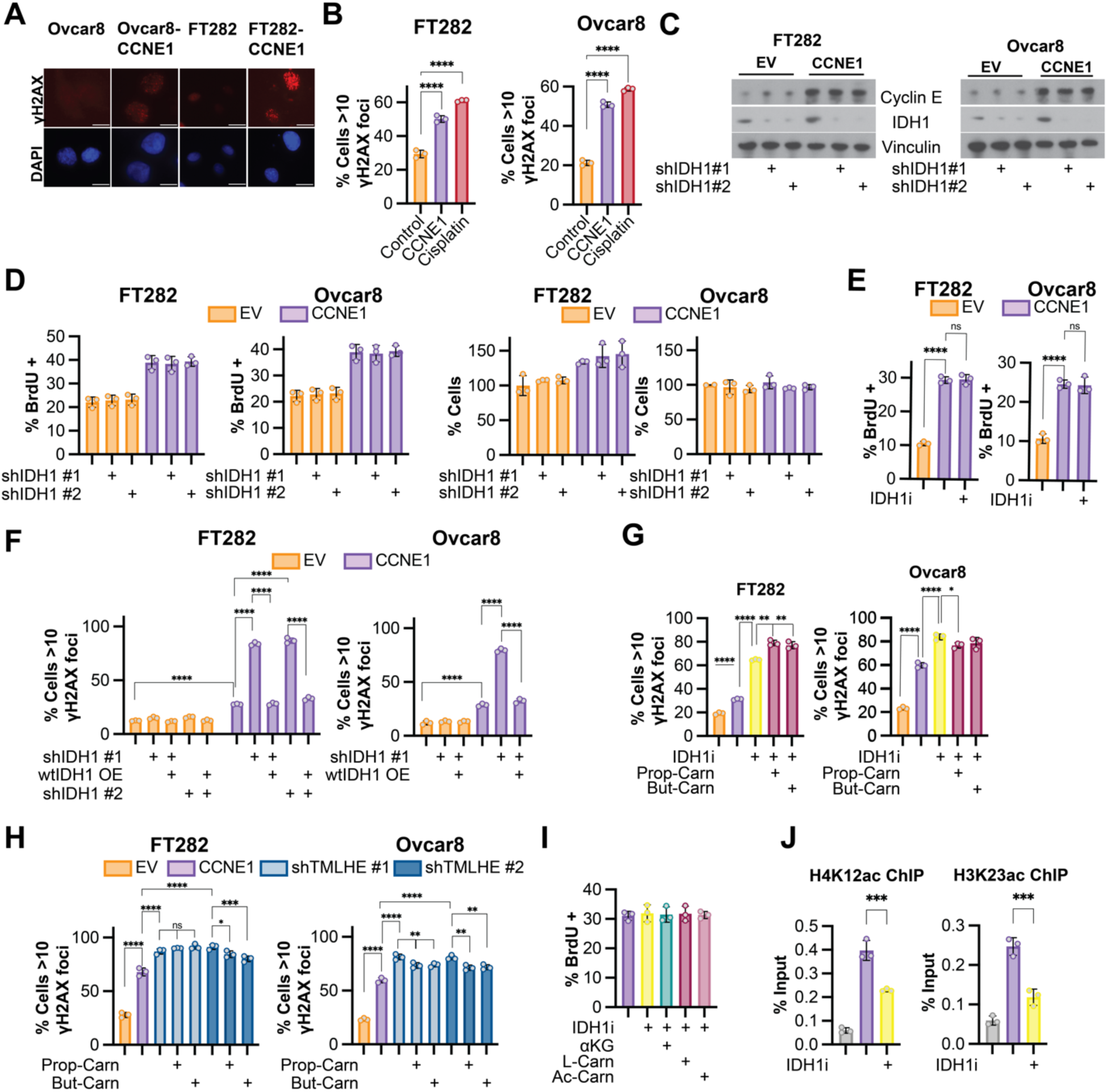
Knockdown or inhibition of IDH1 alone has no effect on proliferation at timepoints associated with increased DNA damage; propionyl-carnitine or butyryl-carnitine do not rescue DNA damage in CCNE1-driven models. Related to Figure 4. **A-B)** Immunofluorescence for γH2AX foci was performed in the indicated isogenic cell lines. **A)** Fluorescent images of DNA damage foci. **B)** Quantification of γH2AX foci. Cisplatin was used as a positive control. **C)** The indicated cells were transduced with two independent shRNAs targeting IDH1 (shIDH1 #1 and shIDH1 #2). Immunoblot analysis of IDH1 was performed. Vinculin was used as a loading control. **D)** The indicated cells were transduced with two independent hairpins targeting IDH1, and BrdU incorporation (left) and % cells by crystal violet staining (right) were assessed. Empty vector (EV) = orange. CCNE1 = purple. **E)** The indicated cells were treated with the IDH1 inhibitor GSK864 (IDH1i), and BrdU incorporation was assessed. Empty vector (EV) = orange. CCNE1 = purple. **F)** The indicated cells were transduced with shRNA targeting IDH1 alone or also transduced with a plasmid overexpressing wildtype IDH1 (wtIDH1). Quantification of γH2AX foci. Empty vector (EV) = orange. CCNE1 = purple. **G)** The indicated cells (EV = orange, CCNE1 = purple) were treated with the IDH1 inhibitor GSK864 (IDH1i) alone (yellow) and supplemented with propionyl-carnitine (Prop-Carn, maroon) or butyryl-carnitine (But-Carn, maroon). **H)** The indicated cells (EV = orange, CCNE1 = purple) transduced with two independent shRNAs targeting TMLHE alone (shTMLHE #1-light blue, shTMLHE #2-dark blue) or supplemented with propionylcarnitine (Prop-Carn) or butyryl-carnitine (But-Carn). Quantification of γH2AX foci. **I)** Quantification of BrdU incorporation of the indicated cells. **J)** U2OS-mCherry-LacI-Fok1 cells were treated with the IDH1 inhibitor GSK864 (IDH1i), and chromatin immunoprecipitation (ChIP) was performed with H4K12ac or H3K23ac antibodies. Shown are representative data from 2-3 independent experiments in each cell line. All graphs represent mean ± SD. *p<0.05; **p<0.01; ***p<0.005; ****p<0.001; ns = not significant

## MATERIALS AND METHODS

### Cells and culture conditions

Ovcar8 cells were a gift from Dr. Benjamin Bitler (University of Colorado). Ovcar8 cells were cultured in RPMI1640 medium (Gibco, cat#11875119) supplemented with 5% Fetal Bovine Serum (BioWest, cat# S1620) and 1% Penicillin/Streptomycin (Fischer Scientific, cat#15-140-122). Ovcar8-Empty Vector (EV) and Cvcar8-CCNE1 cell lines were created by transduction of a Cyclin E1-overexpression plasmid generated by Twist Bioscience (pTwist Lenti SFFV Puro). These cell lines were cultured in RPMI1640 medium (Gibco cat#11875119) supplemented with 5% Fetal Bovine Serum (BioWest, cat#S1620), 1% Penicillin/Streptomycin (Fischer Scientific, cat#15-140-122) and 1ug/ml Puromycin (Gibco, cat#A1113802). FT282-EV and FT282-CCNE1 cells were a gift from Dr. Ronny Drapkin (University of Pennsylvania). FT282 cells were cultured in DMEM:F12 supplemented with 2% FBS and 1% penicillin/streptomycin under 2% oxygen conditions. U2OS-mCherry-LacI-Fok1 cell line was a kind gift from Dr. Roger Greenburg (University of Pennsylvania, Philadelphia, PA) and were maintained in DMEM (Invitrogen, cat#11885084) with 10% FBS and 1% penicillin/streptomycin (Tang et al., 2013). To induce DSBs, cells were treated with Z-(4)-Hydroxytamoxifen (Millipore Sigma, Cat#H7904, 1μM) and Shield-1 (TaKaRa, Cat#632189, 1μM) for a period of 4 hours in charcoal stripped FBS containing medium. HEK293FT cells were used for lentiviral packaging and were cultured in DMEM (Corning, cat#10-013-CV) supplemented with 10% FBS according to ATCC. All cell lines were tested monthly for mycoplasma as described (Uphoff and Drexler, 2005).

### Plasmids, Antibodies, Inhibitors, and Metabolites

All shRNAs were obtained from Sigma-Aldrich. The TCRN are as follows: shIDH1 #1: TRCN0000027253; shIDH1 #2: TRCN0000027249; shTMLHE#1: TRCN0000064804; shTMLHE#2: TRCN0000064807. The pLKO.1 shGFP control was obtained from Addgene (cat#30323). Wildtype IDH1 with overexpression plasmid 3X HA tag was generated by Twist Bioscience (pTwist Lenti SFFV Puro). The following antibodies were obtained from the indicated suppliers: rabbit anti-IDH1 (Cell Signaling Technology, Cat#8137S, 1:1000), rabbit anti-TMLHE (Proteintech, Cat#16621-1-AP, 1:1000), rabbit anti-Cyclin E (Cell Signaling Technology, Cat#4129S, 1:1000), mouse anti-Vinculin (Sigma-Aldrich, Cat#V9131, 1:1000), rabbit pan-acetyl H3 (Active Motif, Cat#61638, 1:1000), rabbit Total Histone H3 (Millipore, Cat#05-928, 1:1000), rabbit pan-acetyl H4 (Active Motif, Cat#39925, 1:1000), rabbit Total Histone H4 (Cell Signaling Technology, Cat#13919S, 1:1000), rabbit H4K8ac (GeneTex, Cat#GTX128957, 1:1000), rabbit H4K12ac (Cell Signaling Technology, Cat#13944S, 1:1000), rabbit H4K23ac (Active Motif, Cat#39131, 1:1000), mouse anti-Beta Actin (Sigma-Aldrich, Cat#A1978, 1:1000), rat anti-BrdU (Abcam, Cat#ab6326, 1:500), mouse anti-phospho-Histone H2A.X (Ser139, 1:500) (Millipore Sigma, Cat#05-636), rabbit anti-53BP1 (Bethyl, Cat # A300-272A, 1:500), mouse BRCA1 (Santa Cruz biotechnology, Cat#sc-6954, 1:500). Secondary antibodies: Fluorescein donkey anti-rat IgG (Jackson ImmunoResearch, Cat#712-095-150, 1:5000), Fluorescein donkey anti-mouse IgG (Jackson ImmunoResearch, Cat#715-095-150, 1:5000), Anti-mouse IgG, HRP-linked (Cell Signaling Technology, Cat#7076, 1:5000), Anti-Rabbit IgG, HRP-linked (Cell Signaling Technology, 7074, 1:5000), Normal mouse IgG (Santa Cruz Biotechnology, Cat#sc-2025, 2μg), Normal rabbit IgG (Cell Signaling Technology, Cat#2729S, 2μg), Fluorescein (FITC)-AffiniPure Donkey Anti-Rabbit (Jackson Immuno, cat# 711-095-152, 1:5000), Cy3 Goat Anti-Rabbit (Jackson Immuno, cat# 111-165-003, 1:5000), Cy3-AffiniPure Donkey Anti-Mouse (Jackson Immuno, cat# 715-165-150, 1:5000). The inhibitors used in this study are as follows: GSK864 (IDH1 inhibitor – MedChem Express, Cat# HY-19540), Olaparib (PARP inhibitor – ApexBio, Cat# A4154), Cisplatin (Selleck Chemicals, Cat# S1166), Mildronate (carnitine synthesis inhibitor – Selleck Chemicals, Cat# S4130), (Z)-4-Hydroxytamoxifen (Millipore Sigma, Cat# H7904), Shield1 (TaKaRa, Cat# 632189). The metabolites used in this study are as follows: αKG (Dimethyl-2-oxoglutarate, Sigma Aldrich, Cat#340631), Triethyl citrate (Sigma Aldrich, Cat#14849), Diethyl succinate (Sigma Aldrich, Cat#112402), L-carnitine hydrochloride (Millipore sigma, Cat#C0283), O-Acetyl-L-carnitine hydrochloride (Sigma Aldrich, Cat#A6706), Propionyl-L-carnitine (Cayman Chemical Company, Cat#9001873), Butyryl-L-carnitine (Cayman Chemical Company, Cat#26542), and N-Acetyl-L-Cysteine (Sigma Aldrich, Cat#A7250).

### Metabolite Measurement

Metabolites were measured by liquid chromatography-high resolution mass spectrometry adapted from previously published approaches (Petrova et al., 2021). Samples were quenched in 1 mL pre-chilled −80°C 80/20 methanol:water (v/v) and spiked with 50µL 1µM isotope labeled TCA cycle mix (Cambridge Isotope Laboratories MSK-TCA-A) pre-diluted in 80/20 methanol:water and 50µL 0.02 ng/µL Propionyl-L-carnitine-(N-methyl-d3) (Sigma Aldrich 52941). After vortexing for 1 min samples were retuned to −80°C for 30 min, centrifuged at 18,000 x g 10 min at 4°C, and the supernatant was transferred to a deep well 96-well plate and evaporated to dryness under nitrogen gas. Samples were reconstituted in 100 µL and 2 µL of the sample was injected from a 4 °C autosampler into a ZIC-pHILIC 150 × 2.1 mm 5 µm particle size column (EMD Millipore) with a ZIC-pHILIC 20 x 2.1 guard column in a Vanquish Duo UHPLC System (Thermo Fisher Scientific) at 25 °C. Chromatography conditions were as follows: buffer A was acetonitrile; buffer B was 20 mM ammonium carbonate, 0.1% (v/v) ammonium hydroxide in water without pH adjustment, with a gradient of 0.5 min at 20% A then a linear gradient from 20% to 80% B; 20–20.5 min: from 80% to 20% B; 20.5–28 min: hold at 20% B at a 0.150 mL/min flow rate. Column elute was introduced to a Q Exactive Plus with a HESI II probe operating in polarity switching mode with full scans from 70-1000 m/z with an insource fragmentation energy of 1. Instruments were controlled via XCalibur 4.1 and data was analyzed on Tracefinder 5.1 using a 5ppm window from the predominant ion positive (carnitines) or negative (all other analytes). Area under the curves for each analyte was normalized to the matched internal standard or the nearest surrogate internal standard. For isotope tracing, isotopologue enrichment was calculated using FluxFix (Trefely et al., 2016).

### Crystal Violet Assays

An equal number of cells were seeded in 96 well plates. For IDH1 inhibitor and carnitine synthesis inhibitor studies: cells pre-treated with the IDH1 inhibitor GSK864 (Ovcar8 cells: 10.565μM, FT282 cells: 12.24μM) or mildronate (145μM for both cell lines), and various metabolites such as αKG (330μM), L-carnitine (1mM), O-Acetyl-Carnitine (1mM), Propionyl-L-carnitine (1mM), Butyryl-L-carnitine (1mM), Citrate (333nM), Succinate (1mM), nacetyl-l-cysteine (NAC, 0.5μM) for a period of 5 days, replacing drugs and metabolites every other day. On the fifth day, cells were treated with olaparib (Ovcar8 cells: 4.213μM, FT282 cells: 4.4124μM) or cisplatin (Ovcar8 cells: 0.455μM, FT282 cells: 5.7125μM) every other day. GSK864 and the metabolites were maintained throughout the experiment. Proliferation was assessed at the tenth day by fixing the plates for 5 mins with 1% paraformaldehyde after which they were stained with 0.05% crystal violet. Wells were destained using 10% acetic acid. Absorbance (590nm) was measured using a spectrophotometer (BioTek Epoch Microplate reader). % cells was calculated by normalizing to appropriate controls. For TMLHE knockdown experiments, cells were transduced, selected, and seeded into 96 well plates. Cells were treated with the same doses of drugs as above. For glutamine starvation assays, cells were seeded in the same manner as described above in normal media. The next day, the media was changed to glutamine-free media (Fischer Scientific, Cat#21870076). The cells were treated as above and maintained in glutamine-free media for the duration of the experiment.

### Western blotting

Cells lysates were collected in 1X sample buffer (2% SDS, 10% glycerol, 0.01% bromophenol blue, 62.5mM Tris, pH 6.8, 0.1M DTT) and boiled to 95°C for 10 min. Protein concentration was determined using the Bradford assay (Bio-Rad, cat#5000006). An equal amount of total protein was resolved using SDS-PAGE gels and transferred to nitrocellulose membranes (Fisher Scientific) at 110mA for 2h at 4°C. Membranes were blocked with 5% nonfat milk or 4% BSA in TBS containing 0.1% Tween-20 (TBS-T) for 1h at room temperature. Membranes were incubated overnight at 4°C in primary antibodies in 4% BSA/TBS + 0.025% sodium azide. Membranes were washed 4 times in TBS-T for 5 min at room temperature after which they were incubated with HRP-conjugated secondary antibodies for 1 h at room temperature. After washing 4 times in TBS-T for 5 min at room temperature, proteins were visualized on film after incubation with SuperSignal West Pico PLUS Chemiluminescent Substrate (ThermoFisher, Waltham, MA).

### *In Vivo* Mouse Experiment

Eight week old female athymic nude mice were purchased from Jackson Laboratories (cat# 002109). All mice were maintained in a HEPA-filtered ventilated rack system at the Animal Facility of the Assembly Building of The Hillman Cancer Center at the University of Pittsburgh School of Medicine. Mice were housed up to 5 mice per cage and in a 12-hour light/dark cycle. All experiments with animals were performed in accordance with institutional guidelines approved by the Institutional Animal Care and Use Committee (IACUC) at the University of Pittsburgh School of Medicine. Five million Ovcar8-EV or Ovcar8-CCNE1 ovarian cancer cells were injected intraperitoneally. After allowing tumors to establish for 9 days, mice were randomly assigned to vehicle, olaparib alone, GSK864 alone, or olaparib+GSK864 treatment. For the first week, mice were treated by daily intraperitoneal injection of vehicle or 150mg/kg GSK864 (Cayman Chemical Company, cat#13960) diluted in 100µl of 16.7 (PG):3.3 (DMSO):40 (PEG400):40(H2O). After the first week, GSK864 treatment was reduced to twice a week and maintained at 30mg/kg. Olaparib diluted in 200µl of 6.8(DMSO):60(PEG300):132(H2O) was administered daily by oral gavage for the following 3 weeks. Mice were weighed weekly to assess toxicity. Animals were euthanized at the end of fourth week, and tumor burden was assessed by counting the number of intraperitoneal nodules.

### αKG-dependent dioxygenases CRISPR library construction

We constructed a pooled sgRNA library containing 64 sgRNAs targeting various genes whose enzymes require αKG as a co-factor for their activity in addition to controls targeting intragenic regions as described previously (Joung et al., 2017) (**Table S1**). We used publicly available CRISPR sgRNA design tools that optimize on-target and minimize off-target genome editing (http://crispr.dfci.harvard.edu/SSC/) and pooled human metabolic library (Birsoy et al., 2015) to identify 10 sgRNAs for each gene. The pooled oligo library was synthesized by Twist Bioscience. The oligo library was cloned as previously described (Joung et al., 2017) into lentiCRISPRv2, (Addgene cat#52961). Briefly, the pooled oligo library was amplified using NEB Next High-Fidelity PCR Master Mix (New England Biolabs, cat#M0541S). The PCR product was digested with Esp3I (BsmBI) (Fisher Scientific, cat#FERFD0454) and ligated into the digested lentiCRISPRv2 backbone. The library was sequenced to ensure optimal sgRNA representation achieving 100% coverage.

### CRISPR Drop Out Screen

The human αKG dioxygenases CRISPR KO library containing 64 genes that require αKG as a co-factor for their activity was designed as stated above. The screening was conducted on Ovcar8/Ovcar8-CCNE1 and FT282/FT282-CCNE1 isogenic cells. Briefly, the appropriate number of cells were infected with pooled libraries at an MOI <0.3 to achieve >400-fold library coverage after selection. Selection was conducted with 500μg of Geneticin (Thermo Fischer Scientific, Cat#10131035) for 6 days. Cells were passaged every 2 days and the whole population was seeded in order to maintain the library coverage throughout. After selection, the whole amplified cell population was seeded at a ratio of 500,000 cells/100mm dish and treated with 4 days with DMSO or olaparib (4.213μM for Ovcar8 and 4.413μM for FT282). At the end of the experiments, cells were harvested for genomic DNA extraction using the Zymo Research kit (cat# D4069). sgRNA inserts were PCR amplified using Ex Taq DNA Polymerase (Takara, cat#RR001A) from sufficient genome equivalents of DNA to achieve an average coverage of >200x of the sgRNA library. See primers in Table S8. Pooled PCR amplicons were then sequenced MiSeq V2 50 cycle kit on an Illumina MiSeq sequencer. MAGeCK was used as the bioinformatics pipeline to analyze negatively enriched genes (Li et al., 2014). Data in Fig. 1I represent Log2 fold change of negative score in (CCNE1 + olaparib vs. CCNE1) vs. negative score in (EV + olaparib vs. EV).

### Mass Spectrometry Analysis of Histone Modifications

The cell pellet was resuspended in nuclear isolation buffer [15mM Tris-HCl (pH 7.5), 60mM KCl, 15mM NaCl, 5mM MgCl_2_, 1mM CaCl_2_, 250mM Sucrose, 1mM DTT, 1:100 Halt Protease Inhibitor Cocktail (Thermo Scientific, 78430), and 10mM sodium butyrate]. Nuclei were resuspended in 0.2M H_2_SO_4_ for 1 hour at room temperature and centrifuged at 4000 *x g* for 5 minutes. Histones were precipitated from the supernatant by the addition of TCA at a final concentration of 20% TCA (v/v). Precipitated histones were pelleted at 10,000 *x g* for 5 minutes, washed once with 0.1% HCl in acetone and twice with acetone followed by centrifugation at 14,000 *x g* for 5 minutes. Histones were air dried then resuspended in 10 μL of 0.1 M (NH)_4_HCO_3_ for derivatization and digestion according to (Garcia et al., 2007). Peptides were resuspended in 100 μL 0.1% TFA in water for LC-MS/MS analysis.

Multiple reaction monitoring was performed on a triple quadrupole (QqQ) mass spectrometer (ThermoFisher Scientific TSQ Quantiva) coupled with an UltiMate 3000 Dionex nano-LC system. Peptides were loaded with 0.1% TFA in water at 2.5 µl/minute for 10 minutes onto a trapping column (3 cm × 150 µm, Bischoff ProntoSIL C18-AQ, 3 µm, 200 Å resin) and then separated on a New Objective PicoChip analytical column (10 cm × 75 µm, ProntoSIL C18-AQ, 3 µm, 200 Å resin). Separation of peptides was achieved using solvent A (0.1% formic acid in water) and solvent B (0.1% formic acid in 95% acetonitrile) with the following gradient: 0 to 35% solvent B at a flow rate of 0.30 µl/minute over 45 minutes. The following QqQ settings were used across all analyses: collision gas pressure of 1.5 mTorr; Q1 peak width of 0.7 (FWHM); cycle time of 2 s; skimmer offset of 10 V; electrospray voltage of 2.5 kV. Monitored peptides were selected based on previous reports (Zheng et al., 2012; Zheng et al., 2013).

Raw MS files were imported and analyzed in Skyline software with Savitzky-Golay smoothing (MacLean et al., 2010). Automatic peak assignments from Skyline were manually confirmed. Peptide peak areas from Skyline were used to determine the relative abundance of each histone modification by calculating the peptide peak area for a peptide of interest and dividing by the sum of the peak areas for all peptides with that sequence. The relative abundances were determined based on the mean of three technical replicates with error bars representing the standard deviation.

### Immunofluorescence, BrdU labeling, and DSB foci visualization

Cells were seeded at an equal density on coverslips and fixed with 4% paraformaldehyde. Cells were washed four times with PBS and permeabilized with 0.2% Triton X-100 in PBS for 5min. Cells were blocked for 5 min with 3% BSA/PBS followed by incubation of corresponding primary antibody in 3% BSA/PBS for 1h at room temperature. Cells were washed three times with 1% Triton X-100 in PBS and incubated with secondary antibody in 3% BSA/PBS for 1h at room temperature. Cells were then incubated with 0.15 µg/ml DAPI for 1 min, washed three times with PBS, mounted with fluorescence mounting medium (9 ml of glycerol [BP229-1; Fisher Scientific], 1 ml of 1× PBS, and 10 mg of p-phenylenediamine [PX0730; EMD Chemicals]; pH was adjusted to 8.0–9.0 using carbonate-bicarbonate buffer [0.2 M anhydrous sodium carbonate, 0.2 M sodium bicarbonate]) and sealed. For DNA damage foci, at least 200 cells per coverslip were counted. Cells were considered positive when they contained >10 γH2AX foci.

For U2OS-mCherry-LacI-Fok1 cells, DSBs were induced as described above. Cells were treated with the following inhibitors and metabolites: GSK864 (10μM), αKG (1mM), L-carnitine (1mM) and O-Acetyl-carnitine (1mM) for a period of 5 days, followed by inducing the DSBs with Tamoxifen (1μM) and shield-1 (1μM) for a period of 4 hours. Post 4 hours, the cells were fixed and immunofluorescence was carried out by staining only for DAPI and mCherry DSB foci were counted. For BRCA1-DSB co-localization, IF was conducted in the same manner described above using BRCA1 primary antibody (1:500) and Fluorescein donkey anti-mouse IgG secondary antibody (1:5000). For quantification, BRCA1-mCherry co-localization was assessed by counting their overlap in >150 cells per coverslip.

For BrdU, cells on coverslips were incubated with 1 μmol/L BrdU for 30 minutes. Cells were fixed with 4% paraformaldehyde (PFA), permeabilized with 0.2% Triton X-100, and then postfixed with 1% PFA + 0.01% Tween-20. Coverslips were DNaseI treated for 10 minutes, and the DNaseI reaction was stopped using 20 mmol/L EDTA. Coverslips were then blocked with 3% BSA/PBS and incubated in anti-BrdU primary antibody (1:500) followed by incubation in FITC anti-Rat secondary antibody (1:1,000). Finally, coverslips were incubated with 0.15 μg/mL DAPI, mounted, and sealed. Images were obtained at room temperature using a Nikon ECLIPSE Ti2 microscope with a 20×/0.17 objective (Nikon DIC N2 Plan Apo) equipped. Images were acquired using NIS-Elements AR software and processed using ImageJ.

### Double-strand break (DSB) assay in *Xenopus* egg extracts

DSB reactions were performed as described previously (Barrows et al., 2022). Briefly, 5 ng/μL of plasmid DNA was incubated in a high-speed supernatant (HSS) supplemented with ATP Regeneration Mix (ARM; 6.5 mM phosphocreatine, 0.65 mM ATP, and 1.6 μg/mL creatine phosphokinase) and 10 μM nocodazole at 21°C for 20 min. A nucleoplasmic extract (NPE) supplemented with ARM, 3.5 mM DTT, and [α-^32^P] dATP was then added at a 2:1 ratio and reactions were incubated at 21°C for an additional 45 min. AgeI (0.25 U/μL; Thermo Fisher) was then added to reactions to generate a DSB. Where indicated, reactions were also supplemented with IDH1i (500 μM) or ⍺KG (500 μM). At the indicated time points, 1 μL of reaction was withdrawn, diluted 6-fold in Replication Stop Dye (3.6% SDS, 18 mM EDTA, 90 mM Tris-HCl pH 8, 90 mg/mL Ficoll, and 3.6 mg/mL Bromophenol Blue), incubated with 20 mg proteinase K at 21°C for 16 h, and then resolved by 0.8% agarose gel electrophoresis. Agarose gels were dried and visualized by autoradiography.

### Chromatin immunoprecipitation

Chromatin immunoprecipitation (ChIP) was performed as described previously (Leon et al., 2021). Using the U20S-DSB-reporter cell line, cells were first treated with their respective inhibitors and supplementation of αKG or L-carnitine and then induced for 4 hours with 4-Hydroxytamoxifen and Shield1 for the induction of DSBs (Tang et al., 2013). Cells were kept in charcoal stripped FBS containing DMEM medium for all the treatments. Post induction, the media was removed by vacuum aspiration and cells were washed twice with 20ml of ice-cold PBS. The cells were then incubated with 20ml of serum-free low-glucose DMEM warmed to 20-22°C. Immediately, 556ul of 37% formaldehyde was added to each dish to reach a final concentration of 1%. The dishes were rocked at 10-15rpm for 10mins at 20-22°C. The crosslinking was then quenched by adding 3ml of 1M glycine (dissolved in PBS) to obtain a final concentration of 125 mM glycine. The dishes were rocked at 10-15 rpm for another 10 min at 20-22°C. The formaldehyde-containing DMEM was removed and cells were washed 3 times with 20ml of ice-cold PBS supplemented with 1x complete protease inhibitor cocktail. The cells were then scraped and collected in 10ml of ice-cold PBS supplemented with 1x Complete protease inhibitor using a plastic cell scraper and shifted into a 50ml conical tube on ice. Another 10ml of ice-cold PBS with 1x Complete Protease Inhibitor was added to the same dish to collect residual cells and these were then transferred to the same 50ml conical tube. The cells were spun down at 250g for 5 min at 4°C and supernatant was removed from the cell pellet. Cells were lysed in 1 mL ChIP lysis buffer (50 mmol/L HEPES-KOH, pH 7.5, 140 mmol/L NaCl, 1 mmol/L EDTA, pH 8.0, 1% Triton X-100, and 0.1% deoxycholate with 0.1 mmol/L PMSF and the EDTA-free protease inhibitor cocktail). Samples were incubated on ice for 10 minutes and then centrifuged at 3,000 rpm for 3 minutes at 4°C. The pellet was resuspended in 500 μL lysis buffer 2 (10 mmol/L Tris, pH 8.0, 200 mmol/L NaCl, 1 mmol/L EDTA, and 0.5 mmol/L EGTA with 0.1 mmol/L PMSF and the EDTA-free protease inhibitor cocktail) and incubated at room temperature for 10 minutes. Samples were centrifuged at 3,000 rpm for 5 minutes at 4°C. Next, the pellet was resuspended in 300 μL lysis buffer 3 (10 mmol/L Tris, pH 8.0, 100 mmol/L NaCl, 1 mmol/L EDTA, 0.5 mmol/L EGTA, 0.1% DOC, and 0.5% *N*-lauroylsarcosine with 0.1 mmol/L PMSF and the EDTA-free protease inhibitor cocktail). Cells were sonicated using a Branson Sonifier 250 for four cycles of 10 seconds on 50 seconds off. Next, 30 μL of 10% Triton X-100 was added to each tube, and then samples were centrifuged at maximum speed for 15 minutes at 4°C. Antibody–bead conjugate solution (50 μL) was added to the supernatant, and chromatin was immunoprecipitated overnight on a rotator at 4°C. The antibodies used at a concentration of 2μg as are as follows: mouse anti-phospho-Histone H2A.X (Ser139), rabbit H4K8ac, Normal Rabbit IgG, Normal Mouse IgG. The following washes were performed: ChIP lysis buffer, ChIP lysis buffer + 0.65 mol/L NaCl, wash buffer (10 mmol/L Tris-HCl, pH 8.0, 250 mmol/L LiCl, 0.5% NP-30, 0.5% deoxycholate, and 1 mmol/L EDTA, pH 8.0), and TE (10 mmol/L Tris-HCl, pH 8.0, and 1 mmol/L EDTA, pH 8.0). DNA was eluted with TES (50 mmol/L Tris-HCl, pH 8.0, 10 mmol/L EDTA, pH 8.0, and 1% SDS) for 30 minutes at 65°C. Reversal of cross-linking was performed by incubating samples overnight at 65°C. Proteins were digested using 1 mg/mL proteinase K and incubating at 37°C for 5 hours. Finally, the DNA was purified using the Wizard SV Gel and PCR Clean Up Kit (Promega). Immunoprecipitated DNA was analyzed by qPCR using iTaq Universal SYBR Green Supermix (Bio-Rad) using Primer 4 (Tang et al., 2013): Forward-CCACCTGACGTCTAA-GAAACCAT; Reverse-GATCCCTCGAGGACGAAAGG. Conditions for amplification were: 5 minutes at 95°C, 40 cycles of 95°C for 10 seconds and 30 seconds with 62°C annealing temperature. Percent input was calculated using the following formula: 100*2^(Adjusted input - Ct (IP) where adjusted input = Ct input - 6.664.

### Quantification and statistical analysis

GraphPad Prism version 10.0 was used to perform statistical analysis. *t* Test and one-way ANOVA followed by *post hoc* Tukey HSD tests were applied as appropriate. When indicated, *P* values were adjusted according to Benjamini–Hochberg FDR. The significance level was established at *P* < 0.05. Heatmaps were generated using GraphPad Prism.

## Notes

### Competing Interest Statement

The authors have declared no competing interest.

### Summary of Updates

Figure 3 and S3 are updated.

